# Developmental determinants of male bias in medulloblastoma

**DOI:** 10.64898/2026.03.25.714163

**Authors:** Luca Bianchini, Ruijie Xu, David Filipovic, Patricia Benites Goncalves da Silva, Laura Sieber, Vuslat Akçay, Frederik Arnskötter, Piyush Joshi, Jana Nolle, Taha Soliman, Ran Tao, Alexandra Scheuing, Konstantin Okonechnikov, Alexander Atamian, Marc Zuckermann, Giles W. Robinson, Giorgia Quadrato, Paul A. Northcott, Lena M. Kutscher

**Author notes:** co-corresponding Lead contact: Lena Kutscher. equal contribution (Luca Bianchini, Ruijie Xu, David Filipovic).

## Abstract

Boys experience an overall increased incidence of several childhood cancers, including medulloblastoma, a clinically heterogeneous cerebellar tumor. In subtypes of Group 3 and Group 4 medulloblastoma, males are three times more prevalent than females. As medulloblastoma is suspected to initiate during fetal development, we hypothesized that this sex bias reflects a combination of prenatal, sex-specific developmental processes and somatic alterations. To test these hypotheses, we compiled a large multi-omics dataset from children with medulloblastoma, which revealed sex-specific alterations, including frequent loss of the inactive X chromosome in females with Group 4. Generation of a sex-matched single-cell transcriptome atlas of the developing murine cerebellum enabled investigation of putative developmental factors underlying sex bias. Progenitors giving rise to Group 3/4 subgroups were more abundant, more proliferative, and harbored more open chromatin for recruitment of LMX1A and OTX2, master transcription factors defining Group 3/4 identity. Advanced genetically engineered mouse models and human cerebellar organoids were leveraged to determine whether sexual dimorphism arises from intrinsic or extrinsic factors. These models showed that the XY genotype contributed to the phenotype, but the predominant effect was driven by presence of the male gonadal hormone testosterone. Our findings provide a sex-specific genetic and neurodevelopmental explanation for male bias in an aggressive pediatric brain tumor. Outcomes from this study may inform novel treatment strategies delivered according to sex and are likely to be broadly applicable to other sex-biased malignancies arising in early life.

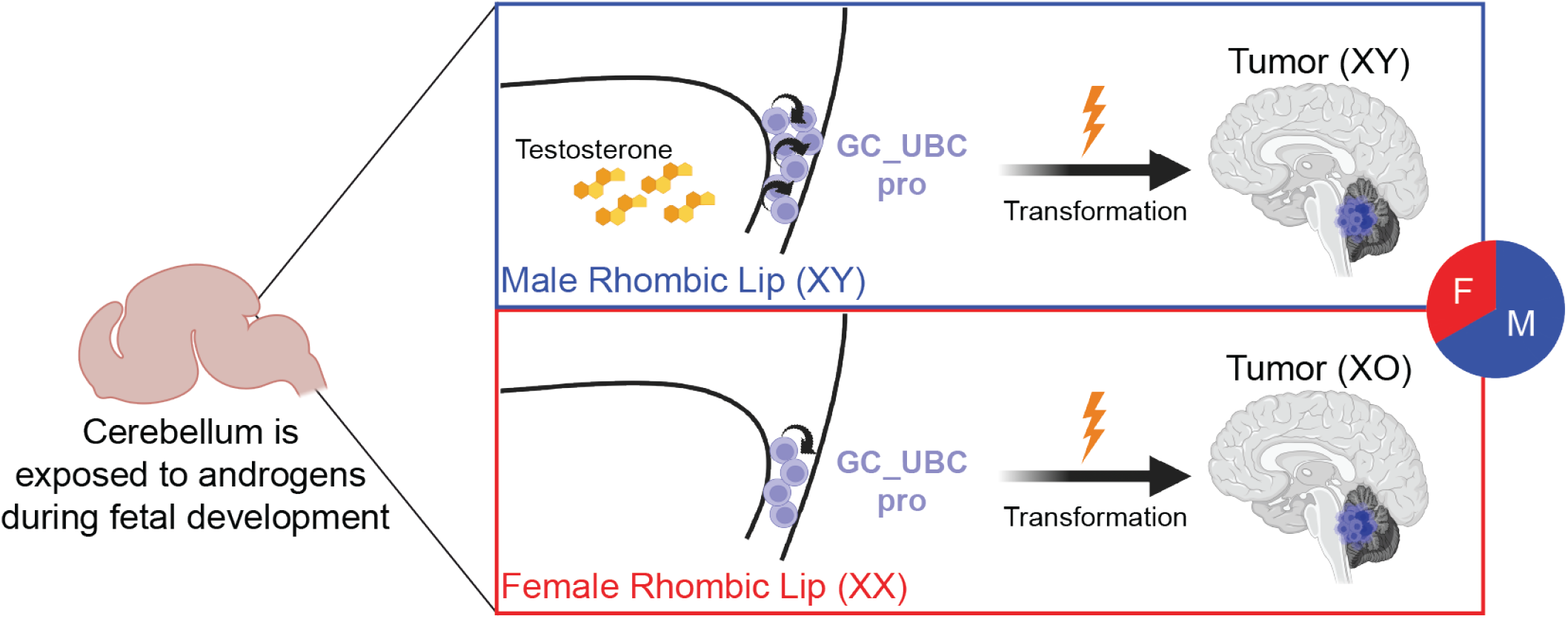

## Introduction

Boys have an overall increased incidence of several childhood cancers across a variety of organ systems, including hematopoietic tumors, sarcomas and brain tumors such as glioblastoma, ependymoma, germinoma, and medulloblastoma (MB)^1,2^. In some cancers, this difference can be partially explained by Escape from X-Inactivation Tumor-Suppressor (EXITS) genes^3^. Namely, X-linked tumor suppressor genes escape X inactivation in female cells and are expressed from both alleles, thus requiring loss of both for complete inactivation. In male cells, these genes are hemizygous and require only a single deleterious mutation to be inactivated^3^. Examples of EXITS genes shown to be more frequently affected by loss-of-function mutations in male tumors include *ATRX, DDX3X, KDM5C*, and *KDM6A*^3^.

MB exhibits a consistent male-biased incidence of approximately 1.5:1^4^. Cellular, molecular, and non-genetic factors contributing to this bias in incidence are largely unknown. This is especially true for Group 3/4 medulloblastoma (G3/4-MB), where biological mechanisms underlying the strong male sex bias have yet to be determined. Prior attempts to uncover the biological basis of the MB sex-bias have yielded inconsistent results^5–8^. Before the molecular era of MB classification, biological sex was reported to be associated with overall survival, with male patients exhibiting poorer survival outcomes^9,10^. However, recent meta-analyses have determined that when accounting for molecular subgroup, sex is no longer independently associated with patient survival^11,12^. This apparent discrepancy is likely explained by the overrepresentation of male patients within the higher-risk G3/4-MBs^13^. These observations suggest that biological sex may preferentially affect tumor initiation or early growth dynamics rather than long-term tumor maintenance.

G3/4-MB comprise a molecularly and clinically heterogenous spectrum of pediatric cerebellar tumors, with a male:female bias of >2:1 that has been consistently reported^14^. A recent pan-cancer whole-genome analysis did not detect sex-biased differences in MB driver mutation prevalence, suggesting that known drivers alone could not explain the sex-biased incidence. However, lack of subgroup stratification in this study likely masked any subgroup-specific differences^15^. Alternatively, these findings suggest that sex-biased incidence could be influenced by developmental factors.Recent developmental studies have determined that these MB subgroups arise from upper rhombic lip-derived progenitors fated towards the unipolar brush cell lineage^16–18^. Initiating genetic events driving G3/4-MB pathogenesis are predicted to occur during fetal development^19^, as early as 12 post-conception weeks (pcw), suggesting that sex-linked differences between males and females may involve factors that are already present before birth. One of the most critical sex-specific processes during fetal development is sex differentiation, which is mediated by the release of gonadal hormones. Beginning at approximately 8 pcw, male gonads start to secrete hormones, including testosterone^20^. Previous studies have shown that testosterone levels peak during critical windows in embryonic and fetal development: embryonic days E16-17 in mouse^21^ and 10-14 pcw in human fetuses^22^. Coincidentally, this developmental window corresponds with the birth and proliferation of the putative cells-of-origin for G3/4-MB, SOX2^+^LMX1A^low^ RL-progenitors (RL^VZ^) and TBR2^+^LMX1A^high^ unipolar brush cell progenitors (GC_UBC progenitors/RL^svz^)^17,18,23^, suggesting that the sex-bias in G3/4-MB may arise from hormone-dependent regulation of these progenitors.

To resolve the biological underpinnings of male sex bias in G3/4-MB, we sought to examine genetic differences according to biological sex in a large multi-omics cohort comprised of >3,000 patient samples. Driver gene alteration frequency and distribution did not profoundly differ between male and female tumors. However, female G4-MBs exhibited frequent loss of the inactive X chromosome, significantly reducing the expression of genes that escape X inactivation, including the chromatin modifier *KDM6A*, a frequently altered gene in this subgroup. In mouse and human cell models, progenitors underlying G3/4-MB development were more abundant and proliferative in the developing male cerebellum in the presence of testosterone. Male progenitors exhibited more accessible binding sites for LMX1A and OTX2 transcription factors, master regulators of G3/4-MB identity. Together, these findings fill a critical gap in our understanding of the developmental determinants of G3/G4-MB and begin to unravel the complex interplay between neurodevelopmental differences in boys and girls that contribute to malignant transformation.

## RESULTS

### Clinical and molecular differences in MB according to sex

To examine associations between biological sex and patient clinical and tumor characteristics, we compared age, histology, metastatic status (M0 or M+), and overall survival between male and female patients across 898 cases from three major North American clinical trials (ACNS0331^24^, ACNS0332,^25^ and SJMB03^26^), separated by clinical trial and stratified by MB subgroup. Our results showed no significant sex-based differences in any of these clinical or tumor characteristics (Figure S1A-D).

To investigate sex-associated molecular differences, we assembled an integrated, multi-omic cohort consisting of 3,026 primary MB tumors, harmonizing genomic (whole-genome and whole-exome sequencing), transcriptomic (expression microarray and RNA-seq), DNA methylation (Illumina 450K or EPIC arrays) data, and clinical annotations from prospective clinical trials^24–27^, retrospective cohorts (International Cancer Genome Consortium, Pediatric Cancer Genome Project, Children’s Brain Tumor Network), and others ^28–30^ (Figure 1A). Most tumors (∼97%) were profiled using DNA methylation arrays, and nearly two-thirds had at least one additional molecular modality available (Figure 1B). The resulting cohort recapitulated the expected distribution of molecular subgroups, dominated by Group 4 tumors; as well as subtypes, with subtypes VII and VIII being most prevalent (Figure 1C). Consistent with prior epidemiological studies, this cohort confirmed a pronounced male bias in overall MB incidence^4^ (1.83 male:female ratio; Figure 1D, S2A). However, stratification by molecular subgroup and subtype revealed a more nuanced and previously underappreciated pattern of sex bias. WNT tumors exhibited a female predominance (0.67 male:female ratio), whereas SHH tumors were male-biased only within childhood and adult-associated subtypes (1.64 male:female ratio in SHH-3 and 2.02 male:female ratio in SHH-4). In contrast, all Group 3 and Group 4 subtypes displayed a strong but variable male bias in incidence (1.77-3.40 male:female ratio; Figure 1D), further emphasizing the importance of molecular resolution when evaluating sex differences in MB.

**Figure 1:**
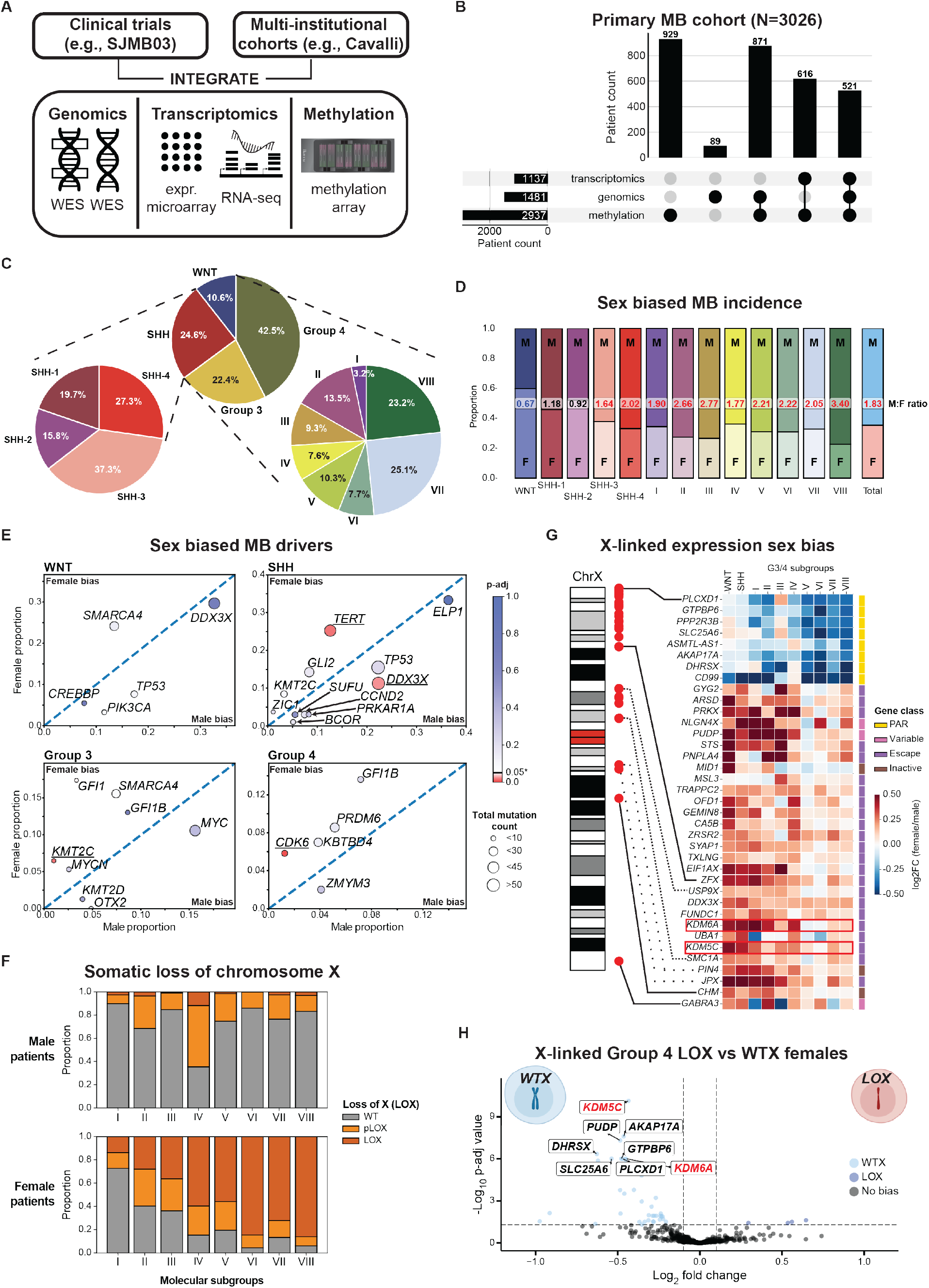
Sex-associated molecular features of primary MB. **A**) Overview of primary medulloblastoma cohort assembly, showing integration and harmonization of genomic (whole-exome sequencing [WES] and whole-genome sequencing [WGS]), transcriptomic (expression microarray and RNA-seq), and DNA methylation (Illumina 450K and EPIC arrays) data from prospective clinical trials (SJMB03, SJYC07, ACNS0331, ACNS0332) and multi-institutional cohorts (Northcott et al.^31^, Cavalli et al.^29^, Capper et al.^28^, Schwalbe et al.^30^). **B**) UpSet plot summarizing the availability and overlap of genomic, transcriptomic, and methylation data across patients in the primary medulloblastoma cohort (N = 3,026). **C**) Distribution of tumors across subgroups and subtypes, including SHH (SHH-1 to SHH-4) and Group 3/4 subtypes (I to VIII). **D**) Sex-associated differences in medulloblastoma incidence across the full cohort, subgroups, and subtype. **E**) Sex-biased somatic driver events stratified by molecular subgroup. Statistical significance was assessed using Fisher’s exact test, comparing the number of detected driver events to the total number of evaluable samples by sex; drivers with p < 0.05 are shown in shades of red and drivers with absolute difference in proportion < 0.02 are not shown. **F**) Proportion of male (top) and female (bottom) patients exhibiting loss of chromosome X (LOX), partial LOX (pLOX), or wild-type X chromosome copy number (WT). **G**) Differential expression analysis of X-linked genes between male and female tumors by subgroup and subtype. **H**) Volcano plot showing differential gene expression between Group 4 female tumors with wild-type X chromosome copy number (WT) and those with LOX. Genes with |log2 fold change| > 0.1 and nominal p < 0.05 are show in light blue (WTX-biased) or gray (LOX-biased). Genes with p < 1E-6 are labelled.

To determine whether sex-biased incidence was accompanied by sex-specific genetic events, we compared the frequency of known/suspected MB driver gene alterations between male and female tumors in a subgroup- and subtype-resolved manner. While a previous pan-cancer analysis reported no sex differences in driver mutation frequencies when primary MBs were analyzed as a single entity^15^, our stratified analyses uncovered subtype-specific sex biases (Figure 1E). Notably, *TERT* promoter mutations in SHH (male:female -12.6% and p-value 0.016), *KMT2C* and *GFI1* alterations in G3 (male:female -5.5%, -14.1% and p-value 0.019, 0.047, respectively) and *CDK6* amplifications in G4 (male:female -4.6% and p-value 0.004) were enriched in females and *DDX3X* alterations in SHH (male:female +11.0% and p-value 0.026) were enriched in males (Figure 1E). Strikingly, loss of chromosome X (LOX) emerged as a highly recurrent and strongly female-biased event across G3/4-MBs, with the highest prevalence observed in G4 subtypes (average LOX=72%; partial LOX (pLOX)=14%) (Figure 1F, S2B-C). Comparative analysis of methylation on chromosome X indicated that LOX in females universally involved the inactive X chromosome (Figure S2D-F).

The observed LOX in females with G3/4-MB suggested a potential link between sex chromosome dosage and subgroup-specific tumor biology. To assess transcriptional differences linked to sex chromosomes, we performed subgroup- and subtype-specific differential expression analyses between male and female tumors. These analyses revealed widespread dysregulation of X-linked genes, with distinct patterns of dysregulation depending on gene class (Figure 1G). Genes located within the pseudoautosomal regions (PARs), which undergo recombination between X and Y chromosomes, generally did not exhibit sex-biased expression; however, pronounced male-biased expression of PAR genes was observed in subtypes VI, VII, and VIII. In contrast, many genes known to escape X chromosome inactivation displayed female-biased expression across WNT and SHH subgroups, a pattern markedly attenuated in G3/4-MB (Figure 1G). Among genes escaping X inactivation, *KDM6A* showed particularly notable dysregulation. *KDM6A* is frequently targeted by loss-of-function mutations in G3/4-MB, subtype VIII^31^, and its reduced expression was consistent with the high prevalence of LOX in female tumors. Together, these findings suggested that loss of the inactive X chromosome reshapes X-linked gene dosage in female MB.

To further examine the consequences of LOX, we stratified female G4 tumors into those with LOX and those retaining both X chromosomes (WTX). Differential expression analysis of X-linked genes revealed a consistent and pronounced reduction in expression of multiple PAR genes (including *AKAP17A, PLCDX1*, and *DHRSX*), as well as the X-linked chromatin regulators *KDM6A* and *KDM5C*, both of which encode lysine-specific demethylases that converge on regulation of H3K27me3 and H3K4me3 chromatin states (Figure 1H).

Collectively, these results establish loss of the inactive X chromosome as a defining and highly recurrent feature of female G4-MB, with direct consequences for the dosage of X-linked chromatin regulators. Although these findings do not provide an immediate explanation for the male bias seen in G3/G4-MB, they suggest that female tumors assigned to these subgroups select for a genotype similar to males, and further implicate deregulation of chromatin modifiers as a prominent theme in the pathogenesis of these tumors^32,33^.

### Increased proliferation in male cerebellar progenitors

We next investigated whether differences in cerebellum development may contribute to sex bias of G3/G4-MB. To that end, we generated a single-cell atlas comprising 320,328 cells and nuclei from sex-matched pre- and post-natal mouse cerebellum, collecting samples at four embryonic days (E12, E14, E16, E17) and at three post-natal days (P0, P2, P4) (Figure 2A-B, S3A). In both sexes, all the expected cell types present at these developmental time points were recovered (Figure S3B), including RL-progenitors (RL-pro), unipolar brush cells (UBC), and UBC / granule cells (GC_UBC) progenitors, granule neuron progenitors (GCPs), granule neurons (GNs), neural stem cells (NSC), astroglia, glutamatergic cerebellar nuclei (Gluta CN), GABAergic cerebellar nuclei (GABA CN), Ventricular zone neuroblast (VZ neuroblast), Interneurons and Purkinje cells (Figure 2B, S3A).

**Figure 2:**
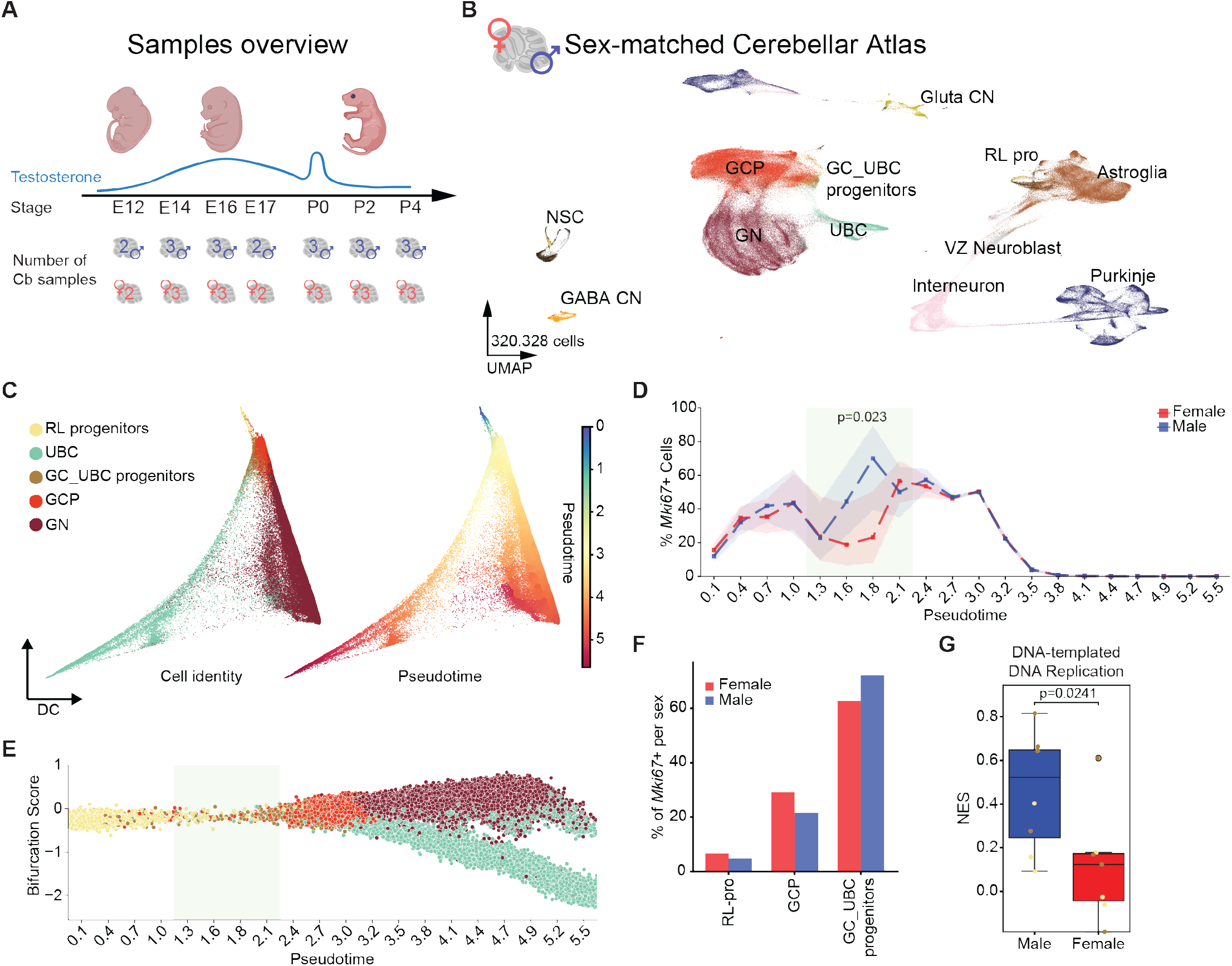
Male murine embryos have more cycling GC_UBC progenitors. **A**) Overview of the sex-matched cerebellar atlas samples. Time points include embryonic day 12, 14, 16, 17, and post-natal days 0, 2, 4. Two or three independent samples have been sequenced per sex per time point. **B**) UMAP showing all the cells recovered in the sex-matched cerebellar atlas. Gluta CN, Glutamatergic cerebellar nuclei. RL pro, Rhombic lip progenitor. GCP, Granule cell progenitor. GC_UBC progenitor, unipolar brush cell and granule cell progenitor. UBC, unipolar brush cell. GN, granule cell. NSC, neural stem cell. GABA CN, GABAergic cerebellar nuclei. VZ Neuroblast, Ventricular zone neuroblast. **C**) Male and female RL-lineage cells (RL-pro, UBC, GC_UBC progenitors, GCP, GN) aligned per pseudotime. Left, cells colored by identity. Right, cells colored by pseudotime. **D**) Percentage of *Mki67*+ cells (normalized expression of *Mki67* > 0) per sex along the pseudotime. Between pseudotime 1.128-2.257, males have more *Mki67*+ cells. Significance assessed using a two-sided t-test. **E**) The pseudotime range in which the male cerebellum has more *Mki67*+ cells corresponds to RL progenitors differentiating into GC_UBC progenitors. **F**) Percentage of *Mki67*+ cells per sex in RL-derived progenitors. Within the GC_UBC progenitor pool, the male cells cycle more (*Mki67*+). **G**) Single sample geneset enrich-ment analysis (ssGSEA) normalized score (NES) of the displayed gene set in pseudobulked male and females RL-pro and GC_UBC progenitors. Significance was assessed using a two-sided t-test.

To investigate sex-specific features pertinent to the G3/ G4-MB lineage-of-origin, we constructed a diffusion map that recapitulated the developmental trajectories and pseudotime of the upper rhombic lip (RL), including RL-pro, GC_UBC progenitors, GCP, GN and UBC (Figure 2C). As expected, male and female cerebellum were similar in the overall cell composition of the excitatory lineage across the analyzed time points (Figure S3C-D). Therefore, we examined the proportion of proliferative cells in the RL lineage marked by *Mki67* expression across developmental pseudotime in male and female mice separately, annotating cells with normalized expression of *Mki67* > 0 as *Mki67*+. This approach revealed a developmental window (pseudotime 1.128-2.257) in which the proportion of *Mki67*+ cells was more pronounced in male than in female cerebella, with a significant difference in abundance between pseudotime 1.692 and 1.975 (Figure 2D). Overlaying mouse age onto developmental pseudotime indicated that this window was predominantly enriched for cells at E16, with cells within this window corresponding to RL-pro differentiating into GC_UBC progenitors (Figure 2E, S3E). Cell-type distribution analysis of *Mki67*+ cells in this developmental window revealed that males had a higher proportion of mitotic GC_UBC progenitors compared to females (Figure 2F), implying that the sex-biased proliferative window is also biased towards this lineage. To determine transcriptional differences between male and female cells, we performed differential expression analysis using pseudo-bulked RL and GC_UBC progenitors from this developmental window. Gene set enrichment analysis identified “DNA-templated DNA replication” as the only significantly upregulated gene set in male compared to female cells (Figure 2G), supporting the higher proliferative state of male GC_UBC progenitors.

Collectively, these results indicate that the G3/G4-MB lineage-of-origin, namely GC_UBC progenitors, is more proliferative and abundant in cerebellum of male mice at E16, motivating deeper analyses of this developmental state.

### Increased abundance of TBR2+ cells at E16 in male cerebellum

G3/G4-MB originate from RL progenitors that express the excitatory lineage marker *LMX1A*^34^. In humans, the RL is subdivided in two domains, the RL ventricular zone (RL^VZ^) and the RL subventricular zone (RL^SVZ^)^23^, with *TBR2*-expressing RL^SVZ^ progenitors thought to represent the developmental origin of G3/G4-MB^17,18^. Although these progenitors were initially proposed to be human-specific, mitotic *TBR2*+ cells (also called *EOMES*+) in the RL have also been identified in mice, where they are referred as GC_UBC progenitors^35,36^. To determine whether the *Tbr2+* GC_UBC lineage is more abundant in male vs female embryos, we crossed *Tbr2-IRES-GFP* reporter mice^37^ with *Lmx1a-Cre*^34^; *LoxP-STOP-LoxP::tdTomato* (Ai9) reporter mice (Figure 3A). On its own, the *Tbr2-IRES-GFP* reporter line labels both *Tbr2+* UBC and *Tbr2+* GNs^38^. The *Lmx1a-Cre* driver labels UBCs and a partially overlapping subset of *Lmx1a*-derived GNs^34^. By intersecting the GFP+ and tdTomato+ cells, we enriched for GC_UBC progenitors at E16 (Figure 3A-B). As *Lmx1a-Cre* resides on chromosome X, the transgene is inactivated in half of the female cells^39^, resulting in only a subset of female *Lmx1a-Cre* lineage cells expressing tdTomato+. However, we detected similar tdTomato expression in male and female RL (Figure 3B).

**Figure 3:**
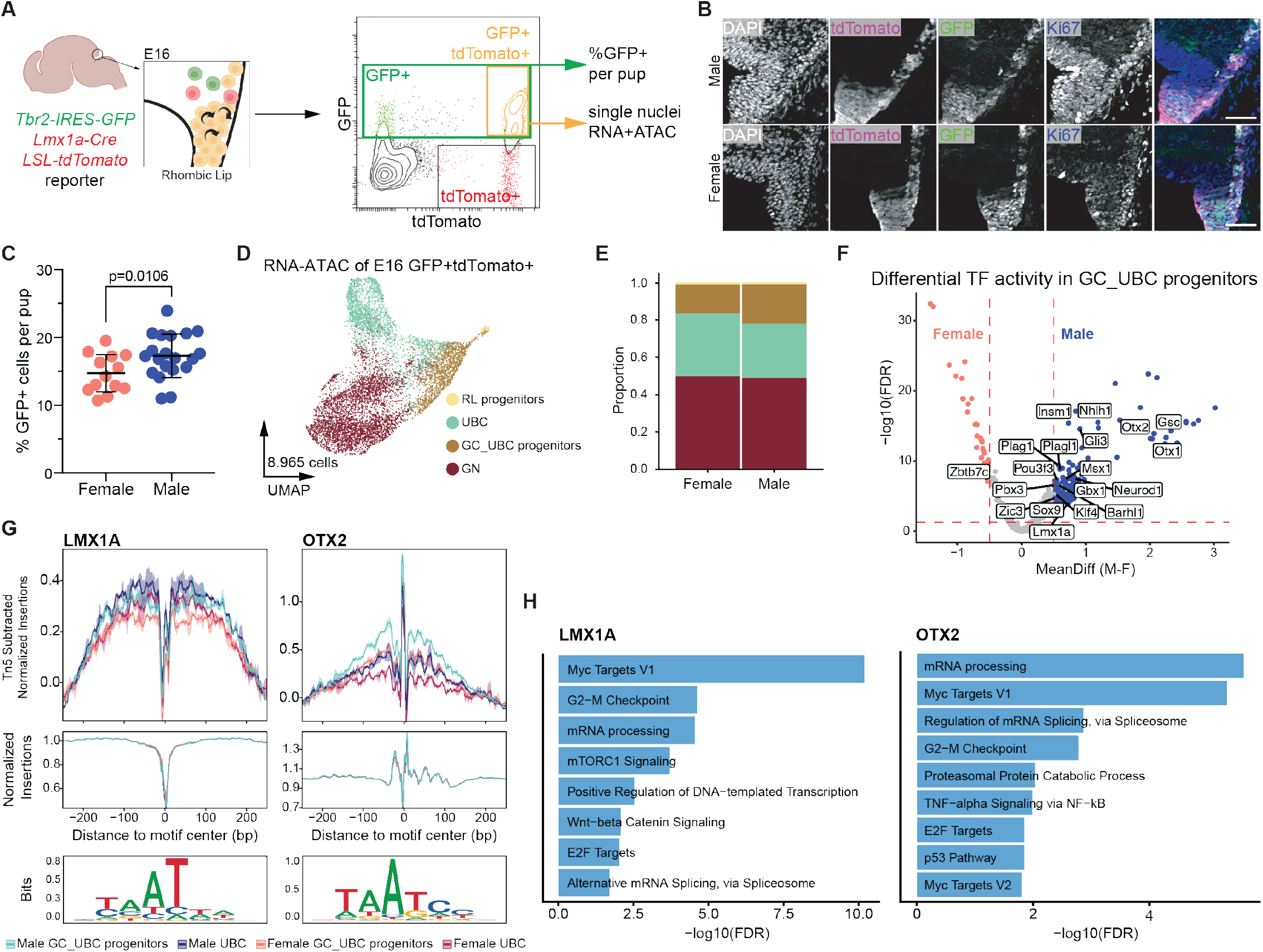
The developing male cerebellum has more Tbr2+ GC_UBC progenitors than the female cerebellum. **A**) The *Tbr2-IRES-GFP; LoxP-STOP-LoxP::tdTomato* (Ai9) reporter mouse line was crossed with the *Lmx1a-Cre* mouse line. The cerebellums of single embryos were dissociated at E16, and the percentage of GFP+ cells were compared between sexes. E16 GFP+/ tdTomato+ cells (enriched for GC_UBC progenitors) were pooled by sex and analyzed by single-cell RNA+ATAC Multiome sequencing. **B**) Representative immunofluorescence of sagittal male and female E16 rhombic lip from *Tbr2-IRES-GFP;Lmx1a-Cre;LoxP-STOP-LoxP::tdTomato* pups. Ki67, blue. TdTomato, magenta. GFP, green. Scale bar, 50 µm. **C**) Percentage of GFP+ cells in male vs female *Tbr2-IRES-GFP;Lmx1a-Cre;LoxP-STOP-LoxP::tdTomato* cerebellum at E16 quantified by flow cytometry. Each dot represents an individual cerebellum, pooled from 5 litters, with mean +/-StdDev shown. Significance determined by unpaired t-test. **D**) UMAP of single-nuclei RNA+ATAC Multiome of E16 GFP+TdTomato+ sorted cerebellar cells. **E**) Proportion of clusters per sex of the Multiome data. Chi square test confirms that males have more UBC cells (FDR 1.59e-05) and GC_UBC progenitors (FDR 3.84e-13), while no significant difference has been observed for the RL_pro (FDR 0.09) and GN (FDR 0.30). **F**) Differential TF activity in GC_UBC progenitors compared between sexes. Male, blue. Female, red. Only TFs with padj <0.05 and absolute TF activity > 0.5 are considered significantly differentially active. TF regulators with positive correlation (padj < 0.05 & correlation > 0.5) between TF expression and their motif activities is labeled (see *Methods* for details). **G**) Tn5-biases normalized footprints for regions surrounding LMX1A and OTX2 TF motifs in GC_UBC progenitors and UBC cells. **H**) Gene set enrichment analysis for target genes associated with motifs of LMX1A and OTX2 that are significantly more accessible (activity_padj < 0.05 & activity_diff > 0) and with higher expression (expr_log2fc > 0) in male cells compared to female cells.

We first determined whether male cerebella have more cells in the *Tbr2*+ lineage compared with females. Cerebella were dissected at E16 and the percentage of GFP+ cells after dissociation was quantified by flow cytometry (Figure 3C, S4A). Male embryos had a small but significant increase in GFP+ cells compared with female embryos (17.25% ± 2.80% in male, 14.66% ± 2.25% in female; Figure 3C), confirming findings from the sex-matched cerebellar atlas (Figure 2D). To identify molecular differences between male and female cells, we performed single-nucleus RNA+ATAC-sequencing (snMultiome) on GFP+tdTomato+ sorted cells at E16 (Figure 3D, S4A-B). We predicted that by pooling multiple cerebella from independent pups and litters, we would still capture the appropriate cell lineage in approximately equal numbers in both male and female pups. Indeed, we recovered the expected RL-derived cell types in both sexes, and the sorting enriched for cells of the UBC lineage (Figure 3D). Male cerebella had more GC_UBC progenitors (chi-square test, FDR 1.59E-05) and fewer UBCs than female cerebella (chi-square test, FDR 3.84E-13), while the GN and RL-pro proportions were similar among both sexes (Figure 3E). Differential gene expression analysis comparing RL-pro and GC_UBC progenitors between sexes, followed by gene set enrichment analysis, determined that male progenitors exhibited higher expression of cell proliferation-related gene sets, and were enriched for gene sets suggesting differential regulation in transcription between sexes (Figure S4C-E). Analysis of sex-biased TF activity in UBC and GC_UBC progenitors showed more open chromatin in DNA regions with motifs of MB-associated TFs^17^, including LMX1A and OTX2, in male cells (Figure 3F-G, S4F). Gene set enrichment analysis on the target genes of these TFs revealed that male embryos were enriched for terms related to increased proliferation and transcriptional regulation (Figure 3H), linking the observed transcriptional differences to possible sex-specific TF activity.

Together, these results indicate that GC_UBC progenitors, the mouse equivalent of G3/G4-MB developmental origin, are more abundant and more proliferative in the male cerebellum. Moreover, GC_UBC progenitors could be more intrinsically predisposed to transformation as binding sites for master TFs responsible for G3/G4-MB identity are more accessible in males.

### Gonadal hormones and sex chromosome complement promote GC_UBC lineage abundance

The differential abundance of the *Tbr2+* GC_UBC lineage in males at E16 could be driven either by exposure to male gonadal hormones or by sex chromosome complement. To determine which of these factors influenced the abundance phenotype, we crossed female *Tbr2-IRES-GFP* reporter animals with Four Core Genotypes (FCG) males. In FCG animals, the sex-determining *Sry* is deleted from the Y chromosome, and overexpressed from a transgene integrated on chr 3^40^, such that the development of testes and ovaries is decoupled from XY and XX sex chromosome complement (Figure 4A). At E16, we quantified the abundance of GFP+ cells in individual cerebella using flow cytometry (Figure 4A-B). Testes-XY embryos had a higher abundance of the *Tbr2-IRES-GFP*+ GC_UBC lineage compared with ovaries-XX embryos. In contrast, ovaries-XY, ovaries-XX, and testes-XX embryos exhibited similar proportions of GFP+ cells (Figure 4B), suggesting that the increased abundance of *Tbr2*+ GC_UBC progenitors depends on both the genotype and the type of gonads present.

**Figure 4:**
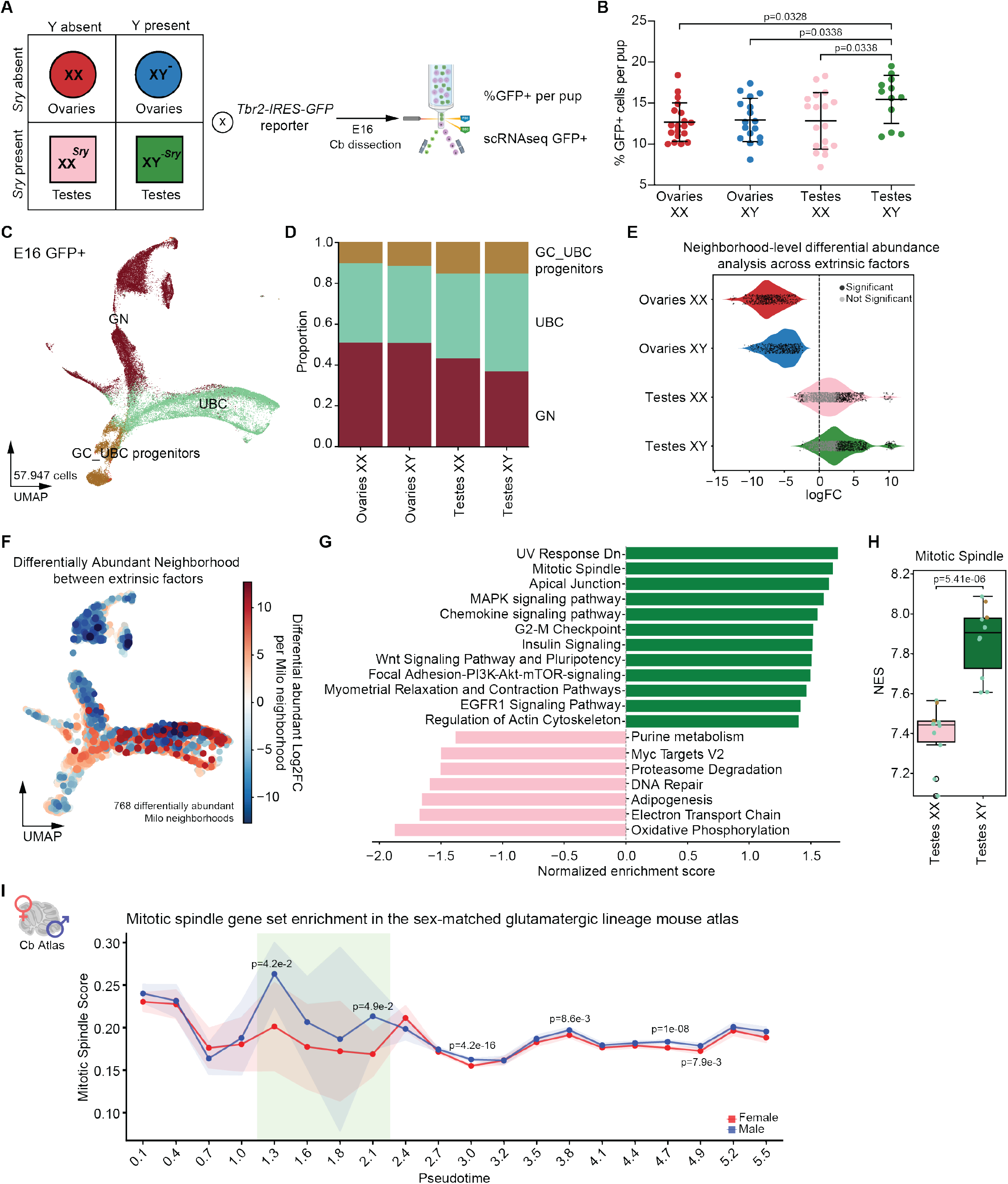
Male gonadal hormones and XY complement promote abundance of GC_UBC progenitors. **A**) Scheme of Four Core Genotypes (FCG) mice, crossed with *Tbr2-IRES-GFP* reporter animals. In FCG animals, sex determining *Sry* gene moved to chromosome 3, decoupling sex chromosome from gonad formation. For analysis, individual cerebellums were dissected and dissociated to single-cells prior to flow cytometry analysis from single pups. **B**) % live GFP+ cells in E16 cerebellum in displayed genotypes. Individual dots, single cerebellum displayed as mean +/-StdDev pooled from 9 litters. Significance determined with one-way ANOVA + Dunnet’s test. **C**) UMAP of single-cell RNA-seq across four core genotypes from *Tbr2-IRES-GFP+* sorted cells. **D**) Proportion of cell types displayed per genotype. **E**) Neighborhood analysis plotted per genotype (<5% SpatialFDR, black true, gray false). **F**) Differential abundance analysis with log_2_-fold change plotted onto UMAP, comparing testes vs ovaries, pooled over sex chromosome. **G**) Enriched gene set pathways in testes XY (green) and testes XX cells (pink). padj < 0.05. **H**) ssGSEA normalized score (NES) of the displayed gene set in pseudobulked testes XY (green) and testes XX cells (pink). Significance was assessed using a two-sided t-test. **I**) Enrichment of the mitotic spindle score in male vs female cells across developmental trajectory.

We next performed single-cell RNA-seq on GFP-sorted cerebellar cells at E16 in each of the four core genotypes to identify putative transcriptional changes underlying these differences (Figure 4C, S5A-C). As expected, we identified cells of the GC_UBC lineage, including GC_UBC progenitors, differentiating UBCs, and differentiating GNs (Figure 4C, S5D). When comparing specific cell types across the four genotypes, a higher proportion of progenitor cells and differentiating UBCs were present in cerebella from embryos with testes, regardless of sex chromosome complement (Figure 4D), suggesting that male gonadal hormones promote differentiation toward the UBC lineage and may influence progenitor cell fate decisions. To test this, we performed differential abundance analysis using Milo^41^, a computational method that identifies neighborhoods of transcriptionally similar cells and tests for differences in their abundance across conditions. Comparing cells from embryos with testes versus ovaries, pooling both XX and XY genotypes, revealed that all significantly abundant cell neighborhoods were driven exclusively by cells from embryos with testes, despite their genotype (Figure 4E). Furthermore, neighborhoods with a significant increase in abundance in the testes-derived samples were highly enriched in the UBC lineage cells (GC_UBC progenitors and UBC), suggesting that male gonadal hormones promote the expansion of the UBC lineage (Figure 4F).

To understand the molecular contribution of the male gonadal hormones to the increase of GFP+ cells in testes-XY mice, we compared testes-XY to ovaries-XY mice using differential gene expression analysis followed by gene-set enrichment analysis (Figure S5E-F). Gene sets associated with stemness were among the statistically significant upregulated pathways in testes-XY embryos, while genes involved in metabolic processes were enriched in ovaries-XY embryos (Figure S5F). As testes-XY embryos had more GFP+ cells than testes-XX embryos, we next examined the role of sex chromosomes in the presence of male gonadal hormones. Testes-XY embryos had higher activation of pathways involved cell cycle and proliferation, while genes involved in metabolic processes were enriched in testes-XX embryos (Figure 4G-H, S5G). Few transcriptional differences were detected when comparing cells derived from embryos with ovaries vs those with testes (ovaries upregulated genes: N=9, testes upregulated genes: N = 4; Figure S5H), suggesting that male gonadal hormones and XY genotype are both required to increase the progenitors’ proliferative status. Interestingly, the chromatin-remodeling genes *Kdm6a* and *Kdm5c* were more highly expressed in XX cells, especially in ovaries-containing animals, highlighting how the loss of the inactive X in G3/G4-MB leads to changes in gene dosage that may contribute to tumorigenesis (Figure S5I).

To further corroborate the proliferative signal identified in the testes-XX vs testes-XY and in the testes-XY vs ovaries-XY comparisons (Figure 4H), we determined a gene score using ssGSEA^42^ across our sex-matched developmental atlas derived from non-transgenic, wild-type outbred animals. We confirmed differential enrichment of the ‘mitotic spindle gene set’ in male cells across development (Figure 4I), including cells within the developmental window that showed a significantly higher proportion of *Mki67*+ UBC cells in male embryos compared to female embryos.

Overall, our findings show that the interaction between male gonadal hormones and XY genotype promote the increased abundance of GC_UBC progenitors in males.

### Testosterone expands the G3/G4-MB lineage-of-origin in human cerebellar organoids

We next sought to determine whether exposure to male gonadal hormone testosterone increases the abundance of RL progenitors in a human cell context. As *LMX1A* is expressed in the RL progenitor zones (RL^VZ^ and RL^SVZ^) from which G3/G4-MBs arise^35^, we generated induced pluripotent stem cell (iPSC) reporter lines expressing mNeonGreen (mNG) fluorescent protein at the 3’ end of the endogenous *LMX1A* gene, in both XX and XY iPSC backgrounds. After confirming that the original and reporter iPSC lines were of high quality (Figure S6A-B, see *Methods* for details), we generated human cerebellar organoids (CbOs) using an established protocol^43^. On day 30 of differentiation, we confirmed that the XX and XY iPSC lines generated CbOs comprising neuronal cerebellar cell types and did not express a forebrain marker, as assessed by immunofluorescence (Figure S6C-D). In addition, we validated that the *LMX1A* reporter lines were of high specificity and sensitivity by quantifying the percentage of cells co-stained with mNG and LMX1A (Figure 5A, S6E). Because mNG expression matched endogenous LMX1A protein expression, we used mNG+ cells as a proxy for LMX1A+ cells.

**Figure 5:**
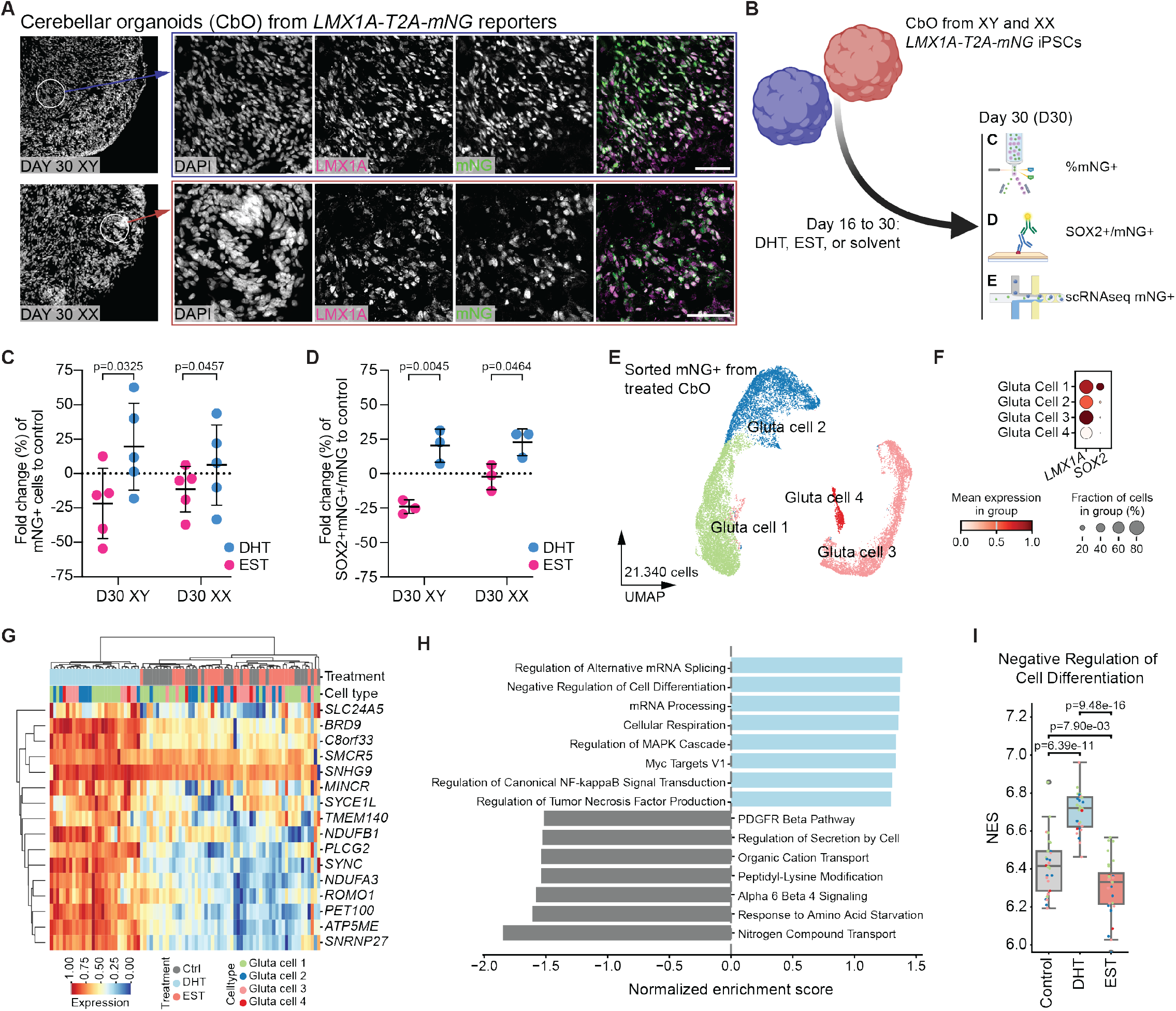
Exposure to testosterone increases LMX1A+ progenitors in human CbOs. **A**) Representative confocal image of cerebellar organoids at day 30 of differentiation and immunofluorescence (IF) of LMX1A (magenta) and mNeonGreen (mNG, green) in day 30 XY and XX *LMX1A-T2A-mNG* reporter human cerebellar organoids (CbO). Scale bar, 50 µm. **B**) Scheme of experimental set-up for gonadal hormone treatment of *LMX1A* reporter CbOs. At day 30 of differentiation, read-outs included %mNG+ cells, IF for SOX2+ cells within the mNG+ population and single-cell RNA sequencing of sorted mNG+ cells across the three treatments. **C**) Flow cytometry measurements of mNG+ (*LMX1A-T2A-mNG* reporter) cells after exposure to estradiol (EST, pink) or dihydrotestosterone (DHT, blue), normalized to solvent control in day 30 XX and XY genetic backgrounds. Displayed as mean +/-StdDev, N = 5 independent experiments, paired t-test. **D**) Percent SOX2^+^mNG^+^ double positive cells relative to total number of mNG+ cells quantified by IF in EST (pink) or DHT (blue) treated day 30 CbOs, relative to solvent control, in XX and XY genetic backgrounds. Displayed as mean +/-StdDev, N = 3 independent quantifications, paired t-test. **E**) UMAP of single-cell RNA-seq across solvent-treated, DHT-treated, and EST-treated sorted *LMX1A-T2A-mNG+* day 30 XX CbO cells. **F**) Relative expression of *LMX1A* and *SOX2* across cell types. **G**) Top differentially expressed genes from DHT/EST treatment vs solvent control. p_adj_ < 0.05. |log_2_ fold change| > 0.5. **H**) Enriched gene set pathways in DHT-treated (blue) and solvent-treated cells (gray). padj < 0.05. **I**) ssGSEA normalized score (NES) of the displayed gene set in solvent-treated (gray), DHT-treated (blue), and EST-treated (red) day 30 organoids. Significance was assessed using a two-sided independent t-test.

To understand the influence of gonadal hormones on LMX1A+ cell populations, we again generated CbOs using the XX and XY reporter cell lines. Following regional specification at day 16 of differentiation, we split the organoid batch into three experimental conditions: treatment with solvent control, with estradiol (EST, estrogen hormone), and with dihydrotestosterone (DHT, a more potent derivative of testosterone, Figure 5B). At day 30 of differentiation (D30), the abundance of mNG+ cells in EST-treated or DHT-treated CbOs was quantified by flow cytometry relative to solvent controls. Treatment with DHT consistently increased the relative abundance of mNG+ cells compared with EST, independent of sex chromosome complement (Figure 5C). As LMX1A^+^SOX2^+^ cells represent progenitors within the RL-UBC lineage, we quantified whether this specific cell type increases with treatment. Consistent with the flow cytometry results, mNG^+^SOX2^+^ cells were more abundant in DHT-treated CbOs compared with EST-treated CbOs, relative to solvent controls, independent of sex chromosome complement (Figure 5D, S6F-G). Interestingly, for both the relative increase in mNG+ cells and the proportion of SOX2+mNG+ among mNG+ cells, the average difference between DHT- and EST-treated CbOs was greater in the XY background (Figure 5C-D). These results are consistent with those obtained with the FCG experiments above, further substantiating that the male gonadal hormone testosterone and the XY genotype cooperate to increase the abundance of specific cerebellar progenitor populations.

To identify transcriptional changes underlying this phenotype, we performed single-cell RNA-seq on mNG-sorted cells at day 30 of differentiation after treatment in XX CbOs. Consistent with broad expression of *LMX1A* in the developing cerebellum, we identified multiple cell types within our CbOs, with only a limited fraction of non-cerebellar cells (Figure 6E, S7A-B). We obtained >6,400 cerebellar cells per condition, and the majority were non-cycling as indicated by the G1 phase (Figure S7C). *LMX1A* expression was highest in glutamatergic neuron (Gluta cell) clusters 1 and 3 (Figure 5F, S7D), and the relative abundance of cell types was similar across the conditions (Figure S7E-F). We performed differential gene expression analysis, comparing gonadal hormone treated cells with solvent control (Figure 5G). DHT treated cells clustered together and away from EST-treated and solvent control cells, which were more similar to each other. Pathways upregulated in DHT-treated cells included “Negative regulation of cell differentiation” and “Regulation of mRNA splicing” (Figure 5H-I, S7G), consistent with a role for male gonadal hormones maintaining the *LMX1A*-derived lineage in a progenitor-like state. Interestingly, DHT-treated CbOs and male murine GC_UBC progenitors both exhibited upregulation of proliferation and mRNA regulation-related pathways, suggesting that human iPSC-derived CbOs recapitulate the biological phenotypes observed *in vivo* (Figure S4E, 5H).

## DISCUSSION

Male predominance amongst children diagnosed with MB is well documented^4,14^. Our detailed assessment of clinical and molecular differences in MB according to sex confirmed a marked male bias in G3/4-MB patients, in agreement with prior studies^13^, while informing how these differences markedly vary among subtypes in the setting of large clinical trials. Our analysis further revealed sex-biased MB drivers and cytogenetics, with pervasive loss of the inactive X chromosome in females with G4-MB. In humans, up to one third of X-linked genes escape X-chromosome inactivation^44^, generally resulting in higher gene dosage of tumor suppressor genes in female cells compared to male^3^. However, since most female G4 MBs undergo LOX, the dosage of these genes becomes balanced between male and female tumors. One such gene is *KDM6A*, which escapes X inactivation and is among the most recurrently altered gene in G4-MB, sustaining inactivating mutations or deletions in ∼10% of cases^31,45–47^. In other male-biased cancers such as T-cell acute lymphoblastic leukemia, *KDM6A* has a significantly higher frequency of deleterious mutations in males^48,49^. However, within subtype VIII MB that exhibits the highest prevalence of LOX in females (86%), *KDM6A* loss-of-function mutations are equally distributed between males and females. Considering the gene dosage impact of losing one copy of X during tumorigenesis, our analysis suggests that the presence of a single X chromosome may influence the incidence rates, providing a partial explanation for the male bias seen in G3/4-MB. Given that large-scale aneuploidy is predicted to be an early event in the majority of G3/4-MB^19^, studies assessing whether (i) genome instability is increased in male fetal cells and (ii) aneuploidy rates are increased after exposure to androgens during development are warranted.

Beyond genetics, our experimental findings suggest that boys are at an increased risk of developing specific subtypes of MB due to an increased abundance of specific progenitor populations in the male cerebellum. Moreover, the chromatin of male progenitors underlying G3/G4-MB is more accessible for LMX1A and OTX2, master regulators of G3/4-MB identity, with OTX2 serving as an essential TF controlling progenitor states^50^. These findings suggest that male cells are intrinsically more prone to transformation, as accessible chromatin persists longer during the relevant developmental window. Indeed, male neuronal progenitors have been previously suggested to be intrinsically more vulnerable to specific driver alterations, including those targeting the chromatin modifier, *Kmt2d*, as deletion of *Kmt2d* in *Nestin*+ cells resulted in a higher MB incidence in male animals^51^. Our *in vivo* results revealed an increase in male cell-of-origin progenitors, which we corroborated in human CbOs. Moreover, CbOs modelling male embryos (XY, DHT-treated) had a higher abundance of cells within the G3/4-MB lineage-of-origin and these cells were maintained in a progenitor-like state. Together, our results derived from both mice and human model systems establish a foundation for future studies investigating sex-specific differences in these progenitors during human fetal development. As organoids successfully mirrored phenotypes observed *in vivo*, they represent a promising platform to dissect molecular mechanisms underlying sex-specific neurodevelopmental processes.

In addition to demonstrating that developing male cerebella have an increased abundance of progenitors underlying G3/4-MB, we provide insights on the molecular determinants underlying this effect. Using the FCG mouse model and genetically engineered iPSC-derived human CbOs, we showed that the XY genotype and the male gonadal hormones interact to promote progenitor expansion. Additional experiments will be required to determine whether the effect is caused by the number of X chromosomes, the modulation of genes on X, or the presence/absence of the Y chromosome^52^. Moreover, other factors are likely to contribute, including sex-dependent regulation of the immune system. Exogenous estrogen has immuno-enhancing effects on humoral immunity^53^, raising the possibility that female brains may also benefit from a more efficient clearance of pre-neoplastic cells.

Although the results obtained from our multifaceted, cross-species studies strongly suggest that the male bias in G3/G4-MB is attributable to a complex interplay among cerebellar progenitor populations, sex chromosome genotype, and the presence of gonadal testosterone, the immediate clinical implications remain uncertain. Androgens have been implicated in other brain tumors, including glioblastoma, where they promote tumor growth and invasiveness. While androgen receptor antagonists have demonstrated potential clinical benefit in the context of adult disease^54–56^, direct antagonism of androgen signaling during development is unlikely to be a viable path forward, given the essential physiological roles this pathway plays in normal growth and sexual determination. However, components of androgen signaling may represent actionable therapeutic targets for males affected by G3/4-MB, motivating a deeper understanding of the mechanisms governing the proproliferative response to testosterone observed in the origins of G3/4-MB. It remains to be determined whether male gonadal hormones and their downstream effectors directly contribute to MB initiation and/or progression. Future studies are needed to establish whether androgen signaling is hijacked to promote tumorigenesis. Furthermore, although primarily recognized for its function in prostate and testis, androgen signaling imposes broad immunomodulatory effects and significantly contributes to sex differences in autoimmunity, infection, and cancer^57^. Future efforts investigating the impact of biological sex and the role of androgens in shaping the G3/G4-MB tumor microenvironment may inform novel immunotherapy strategies specifically engineered for male and female patients.

In summary, we provide an essential foundation towards understanding the dominant male bias observed in children affected by G3/4-MB. We define genetic and developmental determinants of this effect, highlighting the importance of considering sex as an essential biological variable contributing to the pathogenesis of this malignant pediatric brain tumor. Moreover, given the notable male bias observed in various malignancies, our study provides a framework for more broadly resolving sex bias in other cancers arising in early life.

## Data and code availability

Raw data will be provided upon request. Analysis available at github.com/rx32940/MB_XY_bias.

## Acknowledgments

Funding was provided by The Brain Tumour Charity – Quest for Cures consortium grant to L.M.K and P.A.N. (COMMAND). L.M.K. was supported by the Emmy Noether Programme from the German Research Foundation (551030459). P.A.N. acknowledges funding from the American Lebanese Syrian Associated Charities (St. Jude), The Mark Foundation for Cancer Research (Emerging Leader Award), and the National Cancer Institute (1R01CA270785-01A1). We acknowledge the following core facilities at the DKFZ: Genomics core facility, Single-cell open lab, light microscopy core facility, flow cytometry core facility, and the Center for Preclinical Studies. Material was generated from the KOLF2.C1 cell line produced by the Sanger as part of the Human Induced Pluripotent Stem Cell Initiative (HIPSCI) and was generated with the support of the National Institutes of Health (NIH) and the NIH’s iPSC Neurodegenerative Disease Initiative. We thank Esther Becker for sharing AH017.3 iPSC line, Thierry Walzer for sharing the *Tbr2-IRES-GFP* mouse line, Kathleen J. Millen for the *Lmx1a-Cre* mouse line, and James Cleland, Duncan Odom and James Turner for the Four Core Genotypes mouse model. Some figure panels were generated with in BioRender (https://BioRender.com/g6msyuh). We thank Katie Han (St. Jude) for carefully reviewing the manuscript.

## Author Contributions

Human tumor analysis: D.F., K.O.

Single-cell analysis: R.X., L.B., P.J.

Animal experiments: L.B., P.B.G.d.S., L.S., V.A., T.S., L.M.K.

Reporter line generation: L.B., P.B.G.d.S., J.N.

CbO experiments: L.B., F.A., A.S.

Critical data sets: G.W.R.

Critical scientific input: A.A., G.Q., M.Z.

Writing: L.B., D.F., R.X., F.A., P.A.N., L.M.K.

Supervision: P.A.N., L.M.K.

Funding: P.A.N., L.M.K.

---

**Fig. S1:**
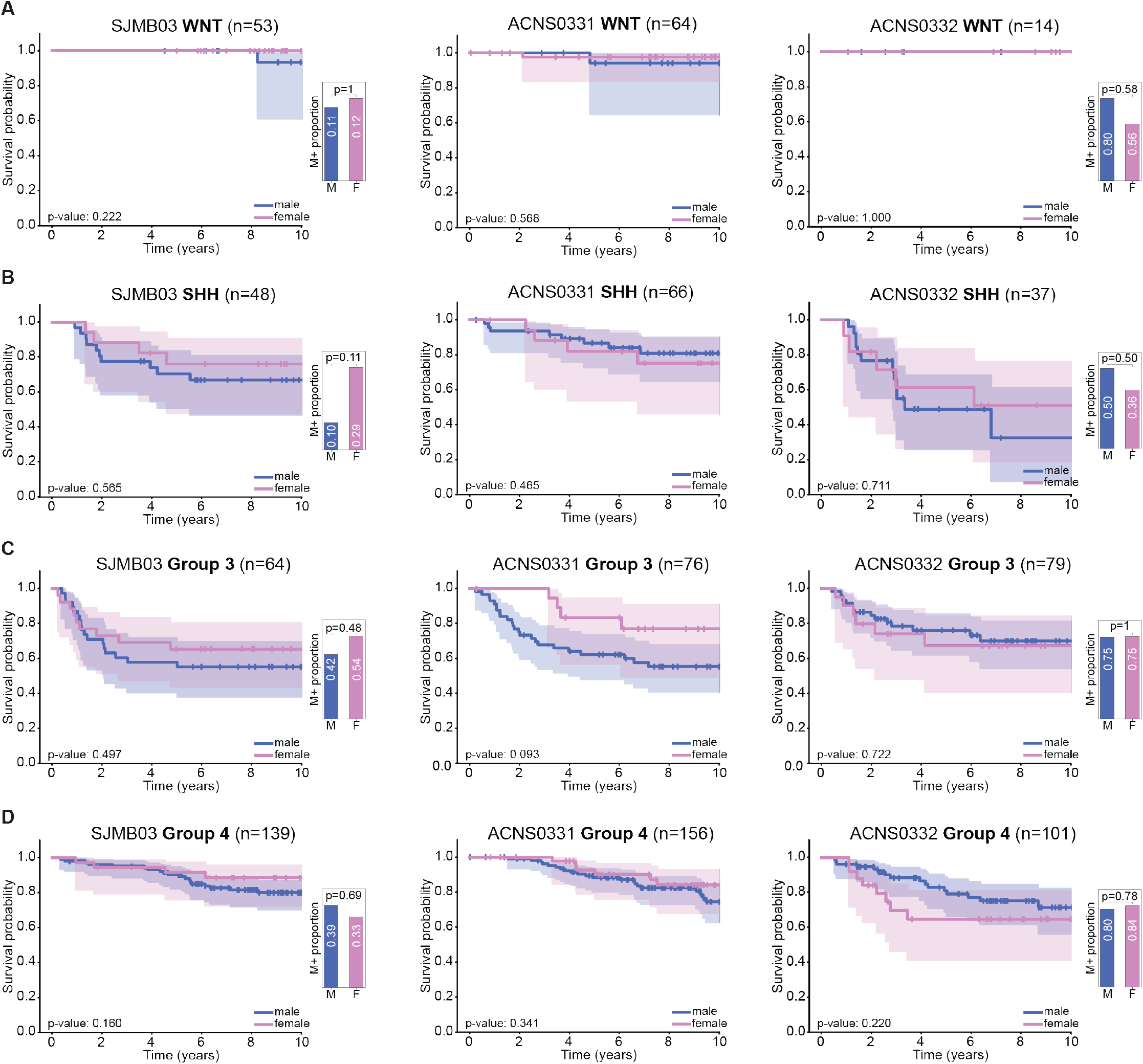
Clinical differences between male and female patients stratified by subgroup and clinical trial. SJMB03 clinical trial (left column), ACNS0331 clinical trial (middle column), and ACNS0332 clinical trial (right column) are analyzed separately. Overall survival curve using Cox regression analysis compared by sex are shown on the left; metastatic status (M0/M+) compared by sex using Fisher’s exact test are shown on the right for each MB subgroup – WNT (**A**), SHH (**B**), Group 3 (**C**), Group 4 (**D**). ACNS0331 had only low risk patients (M0) enrolled.

**Fig. S2:**
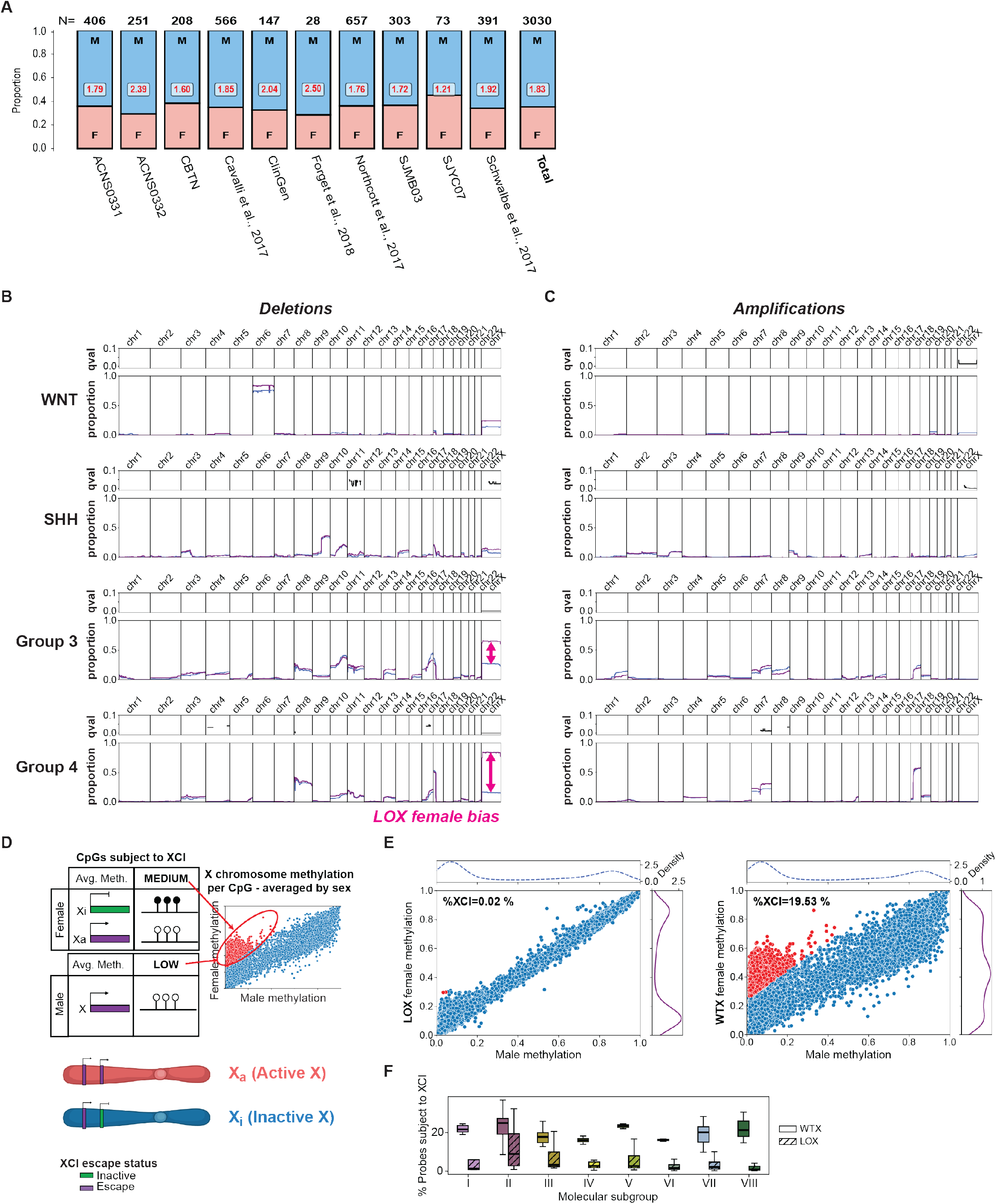
Loss of inactive X in Group 3 and Group 4 MB. **A**) Sex-associated differences in medulloblastoma incidence across each of the trials analyzed. **B**,**C**) Comparison of proportions of genome-wide deletions/amplifications between male and female patients. Each row represents a medulloblastoma subgroup (WNT, SHH, Group 3, and Group 4). Statistically significant deletions/amplifications were identified using Fisher’s exact test with Benjamini– Hochberg correction; q-values are shown as black lines in the upper panel of each row only for significant events (q < 0.05). Proportions of deletions (**B**) and amplifications (**C**) are shown for male (blue lines) and female (purple lines) patients in the lower panel of each row. **D**) Schematic showing the active (Xa) and inactive (Xi) copy of X chromosome in females. Example of a gene subject to X inactivation (XCI) – inactive gene, is shown in green and gene escaping XCI – escape genes, is shown in purple. Schematic illustrating the effect of XCI on average methylation of CpGs in females compared to males. CpGs subject to XCI are methylated at a medium level in females and low level in males. **E**) Example of subtype VIII females with LOX (left) and wildtype X (WTX; right). Each dot represents a single CpG probe on chromosome X, with average male methylation shown on x-axis and average female methylation shown on the y-axis. CpGs with a difference between females and males larger than 0.25 and average male methylation lower than 0.5 are shown in red and are considered as CpGs undergoing XCI. Percent of these probes is shown as %XCI. **F**) Boxplot showing %XCI in WTX (solid fill) and LOX (hatched fill) across G3/4-MB subtypes.

**Fig. S3:**
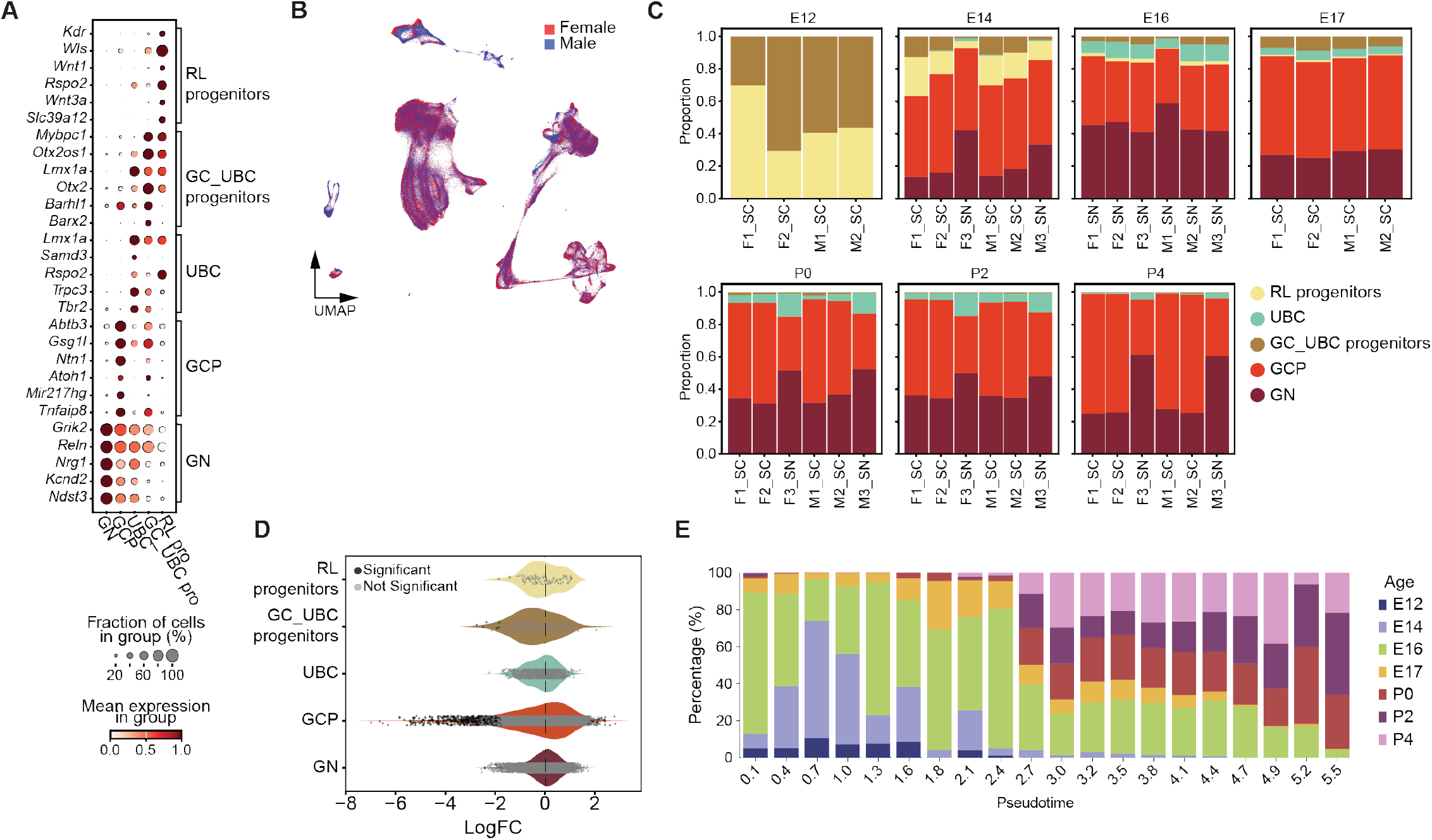
Sex-matched murine cerebellar atlas. **A**) Dot plot of marker genes used for cell annotation of the excitatory lineage. **B**) UMAP showing male and female cells distribution across the dataset. **C**) Proportion per sample per developmental time point of cells deriving from RL progenitors. F, female pup. M, male pup. SC, single cell. SN, single nuclei. **D**) Neighborhood analysis per cluster comparing the relative abundance of male vs female cells (<5% SpatialFDR, black true, gray false). **E**) Sample’s stage distribution along the pseudotime. E16 cerebellar cells are enriched between pseudotime 1.692 and 1.975.

**Fig. S4.**
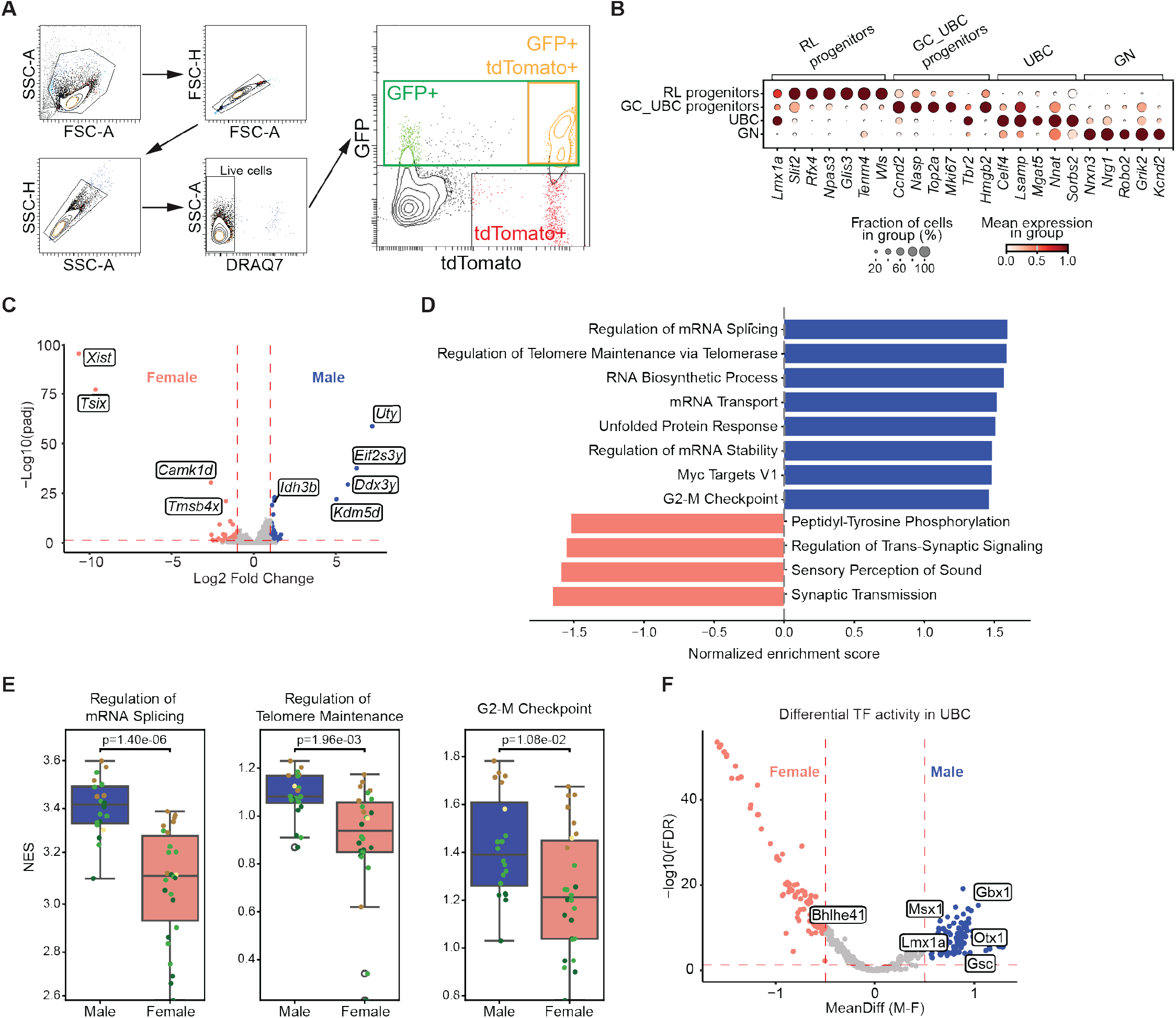
Characterization of the *Tbr2-IRES-GFP; Lmx1a-Cre; LoxP-STOP-LoxP::tdTomato* mouse line. **A**) Gating strategy for sorting single and double positive primary mouse cells. After single cell gating, live (DRAQ7 negative) cells were gated for GFP+ and tdTomato+ expression. **B**) Dot plot of marker genes used for cell annotation. **C**) Differential gene expression analysis comparing male and female RL_pro and GC_UBC progenitors. Only genes with a p_adj_ <0.05 and log_2_ fold change >0.5 considered as differentially regulated. **D**) Enriched gene set pathways in male (blue) and female (red) RL-pro and GC_UBC progenitors. p_adj_ < 0.05. **E**) ssGSEA normalized score (NES) of the displayed gene sets pseudobulked RL-pro and GC_UBC progenitors in male (blue) and female (red). Significance was assessed using a two-sided t-test. **F**) Differential TF activity in GC_UBC progenitor compared between sexes. Male, blue. Female, red. Only TFs with padj <0.05 and absolute TF activity difference > 0.5 are considered significantly differentially active. TF regulators with positive correlation (padj < 0.05 & correlation > 0.5) between TF expression and their motif activities is labeled (see *Methods* for details).

**Fig. S5.**
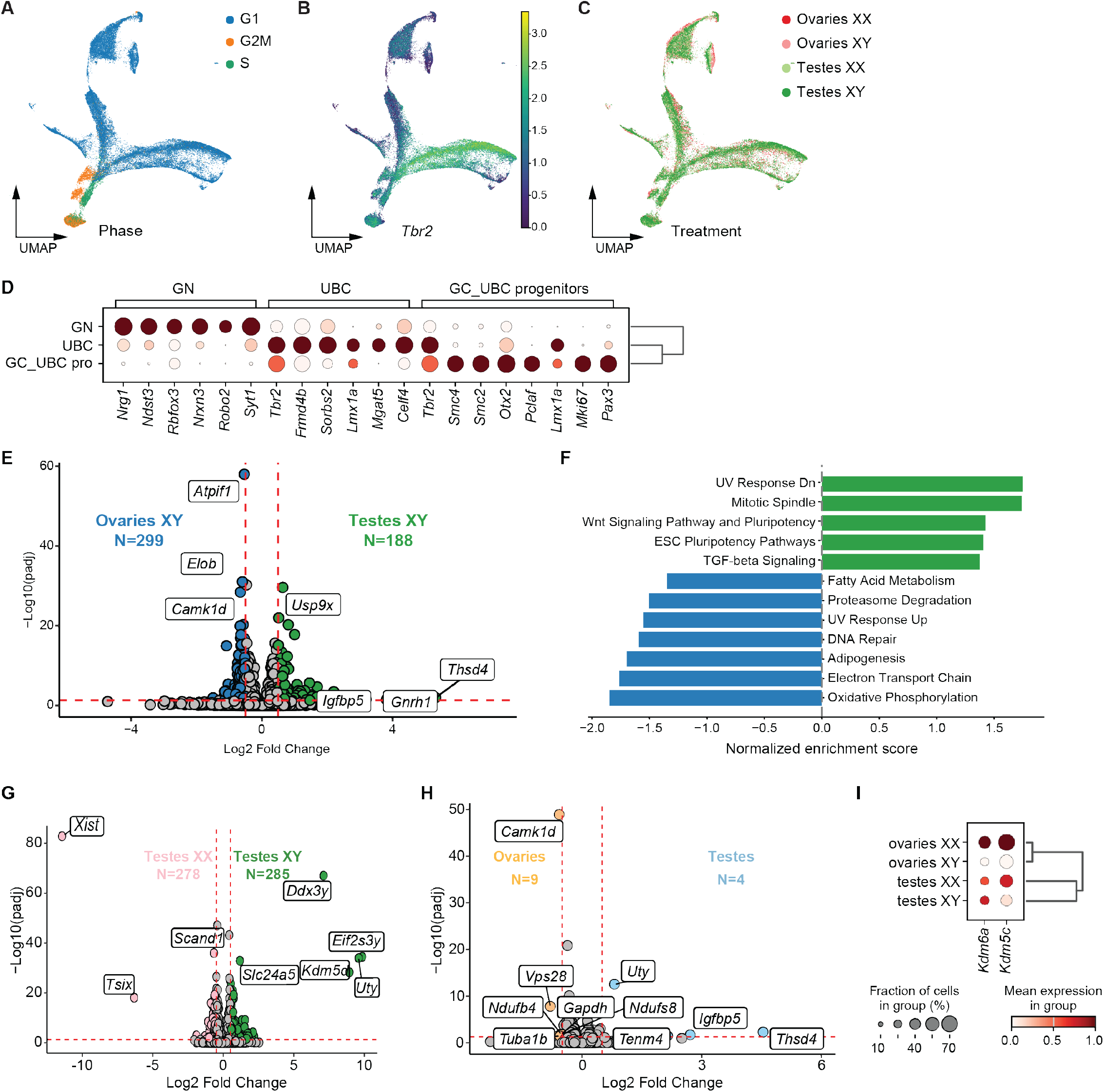
Testes XY cells have an increase in mitotic spindle genes and MAPK signaling pathway genes. **A-C**) UMAP of single-cell RNA-seq across four core genotypes from *Tbr2-IRES-GFP+* sorted cells, colored by A) genotype, B) cell cycle phase, and C) relative *Tbr2* expression. **D**) Dot plot with expression of marker genes for major cell types. **E**) Differential gene expression comparing testes-XY and ovaries-XY animals. Only genes with a p_adj_ <0.05 and log2 fold change >0.5 considered as differentially regulated. N, number of genes meeting these criteria. **F**) Enriched gene set pathways in testes-XY (green) and ovaries-XY (blue). p_adj_ < 0.05. **G**) Differential gene expression comparing testes-XY and testes-XX animals. Only genes with a p_adj_ <0.05 and log_2_fold change >0.5 considered as differentially regulated. N, number of genes meeting these criteria. **H**) Differential gene expression comparing pooled testes and ovaries animals. Only genes with a p_adj_ <0.05 and log_2_ fold change >0.5 considered as differentially regulated. N, number of genes meeting these criteria. **I**) Dot plot with expression of chromatin modifying genes *Kdm6a* and *Kdm5c* in the displayed genotypes.

**Fig. S6.**
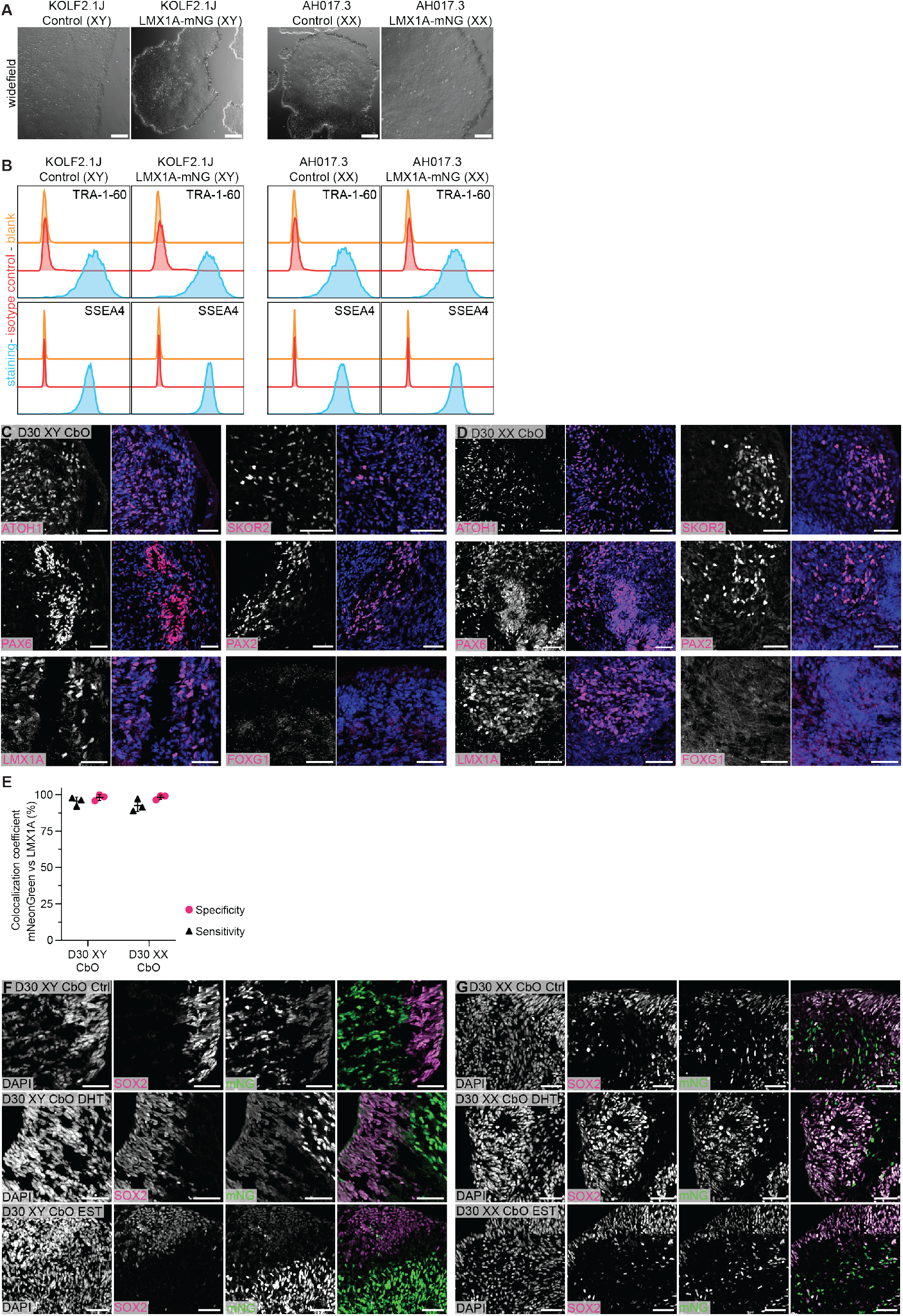
Quality control of iPSC reporter lines and cerebellar organoids (CbO). **A**) Widefield image of control and transgenic iPSC lines. Scale bar, 100 µm. **B**) Expression of pluripotency markers TRA-1-60 and SSEA4 in control and transgenic iPSC lines as quantified by flow cytometry. **C**) Representative IF of day 30 KOLF2.1J-derived cerebellar organoids (D30 XY CbO) for the cerebellar markers ATOH1, PAX6, LMX1A, SKOR2, PAX2 and the forebrain marker FOXG1. Scale bar, 50 µm. **D**) Representative IF of day 30 AH017.3-derived cerebellar organoids (D30 XX CbO) for the cerebellar markers ATOH1, PAX6, LMX1A, SKOR2, PAX2 and the forebrain marker FOXG1. Scale bar, 50 µm. **E**) Quantification of the colocalization between mNG and LMX1A in day 30 XX and XY CbO. Specificity, proportion of LMX1A+mNG+/mNG+. Sensitivity, proportion of mNG+LMX1A+/LMX1A+. **F**) Representative IF of SOX2 (magenta) and mNeonGreen (mNG, green, proxy for LMX1A) in day 30 XY human CbO treated with solvent control, DHT, or estradiol. Scale bar, 50 µm. **G**) Representative IF of SOX2 (magenta) and mNeonGreen (mNG, green, proxy for LMX1A) in day 30 XX human CbO treated with solvent control, DHT, or EST. Scale bar, 50 µm.

**Fig. S7.**
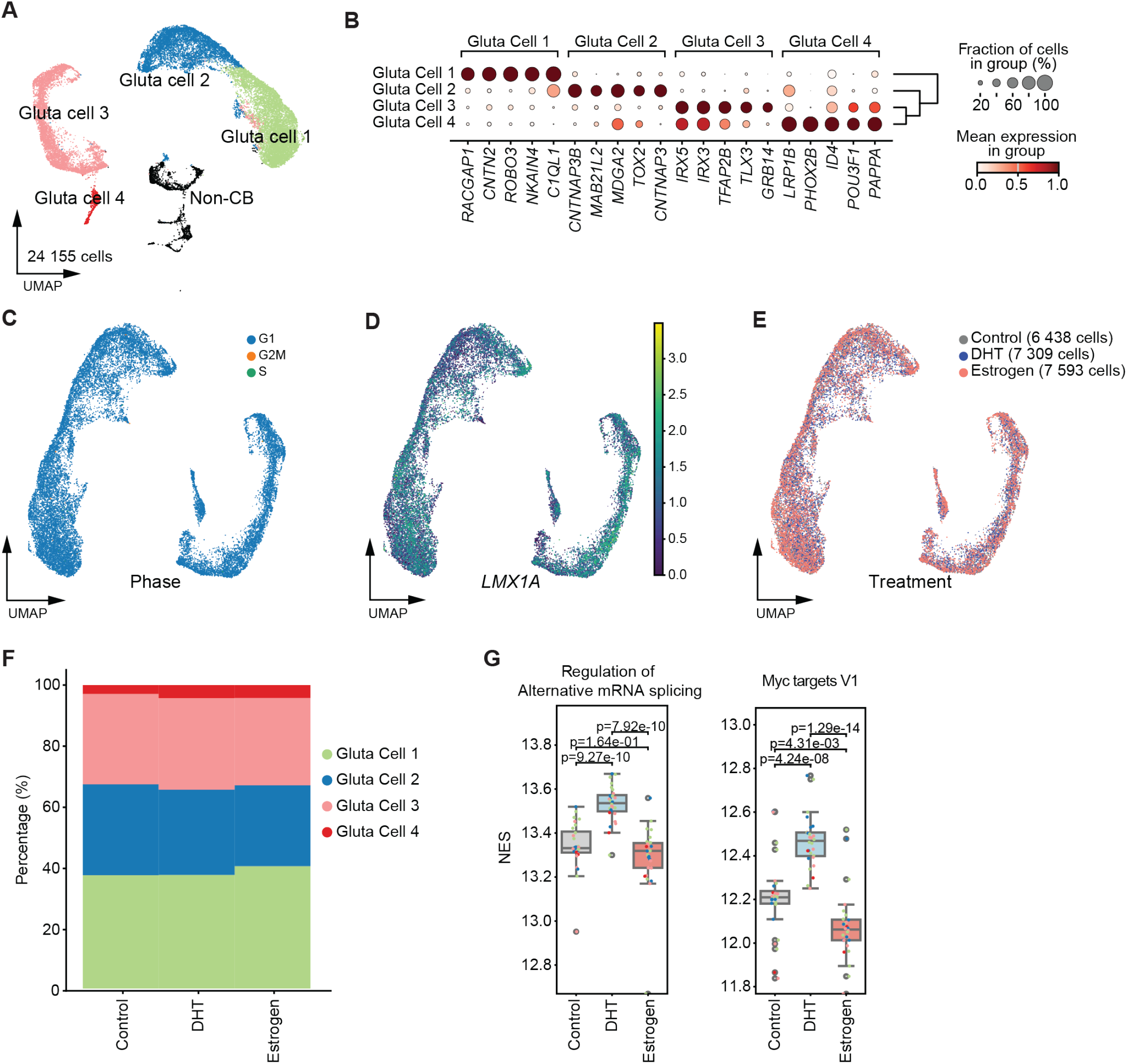
Relative abundance of cell types in treated CbOs. **A**) UMAP of cells recovered from day 30 XX CbOs. Gluta cell, Cerebellar glutamatergic cell. Non-CB, non-cerebellar cells. **B**) Dot plot of marker genes per cluster used for cell annotation. **C-E**) UMAP of single-cell RNA-seq across treated sorted CbO cells, colored by C) cell cycle phase, D) treatment, and E) relative LM1A expression. **F**) Proportion of cell types per displayed genotype. **G**) ssGSEA normalized score (NES) of the displayed gene sets in pseudobulked CbO treated with solvent control, DHT, or estradiol day 30 CbO. Significance was assessed using a two-sided t-test.

## MATERIAL AND METHODS

### Key Resource Table

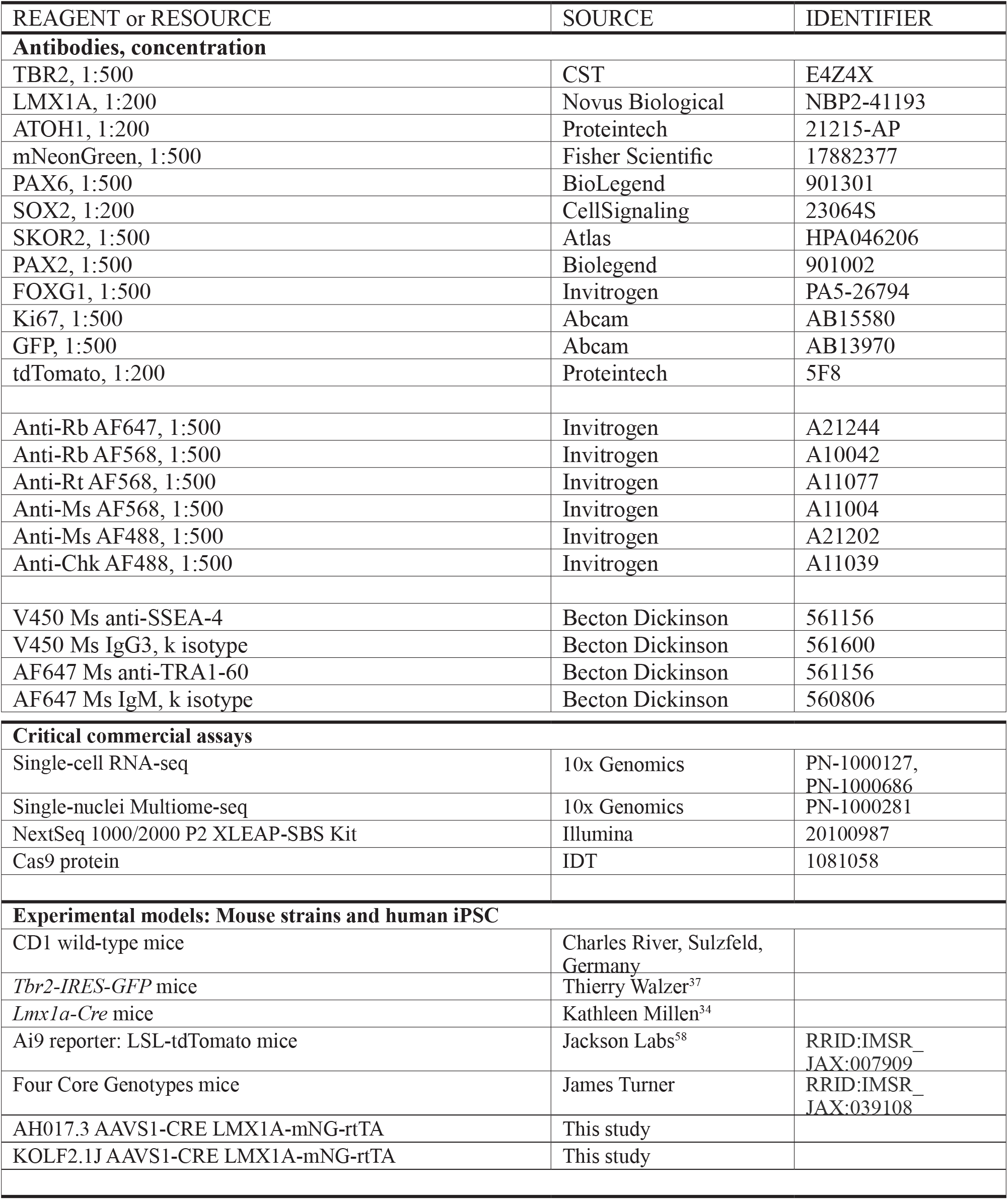
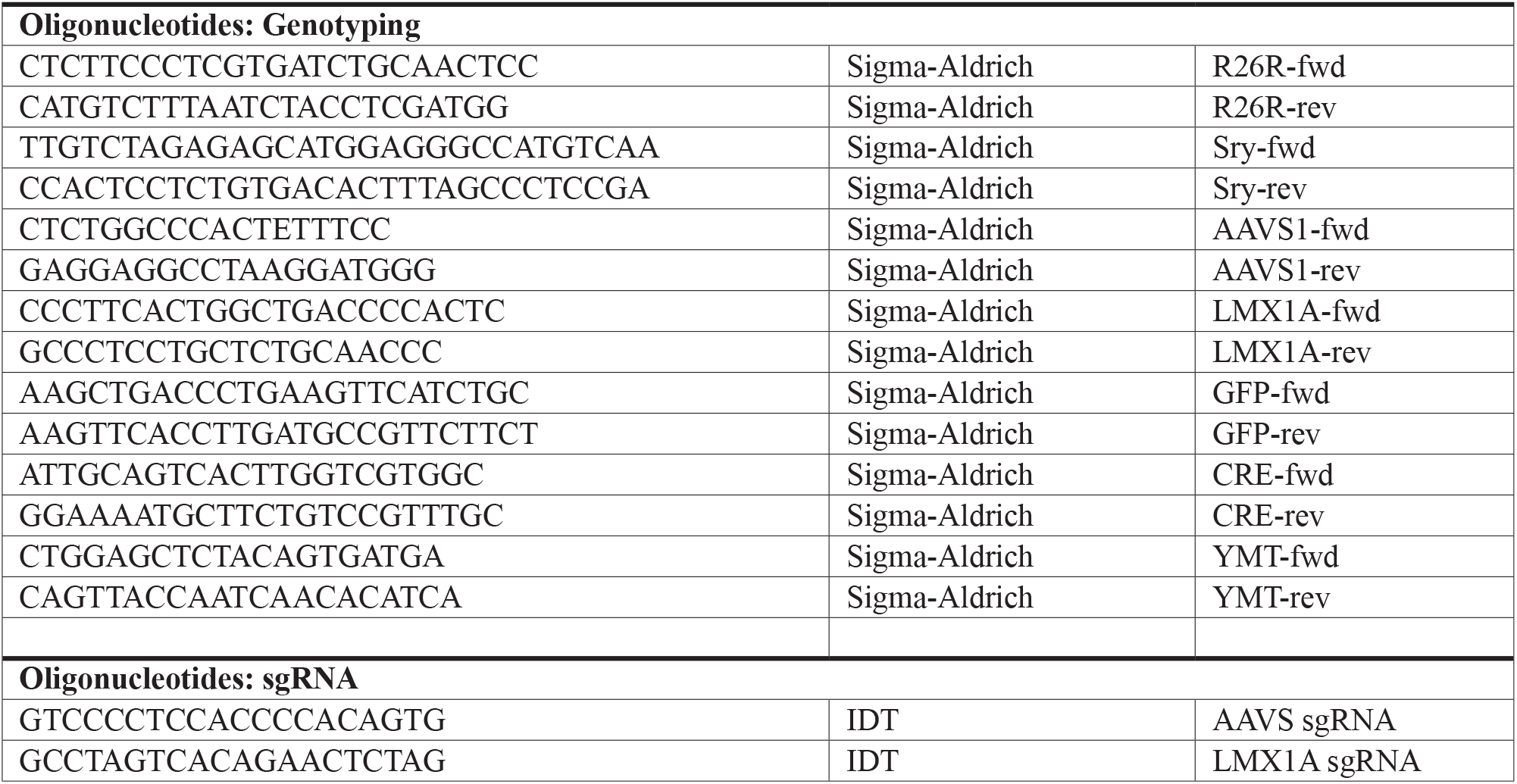

### EXPERIMENTAL MODEL DETAILS

#### Animals

Animal strains used in the study are listed in the Key Resources Table. Animals were housed under a 12h/12h dark/light cycle controlled for temperature (20-24°C) and humidity (40-65%) with *ad libitum* access to food and water. Genetically engineered mice were bred on a C57BL/6N background. Animal experiments for this study were approved by the responsible local authorities: DKFZ Central Animal Laboratory (DKFZ383, DKFZ443-24) and by the Institutional Review Board at St. Jude Children’s Research Hospital (589-100536). The morning of vaginal plug was designated as E0, and the day of birth as P0.

#### Genotyping

Genotyping primer sequences are available in Key Resource Table. Tissue samples were either sent to Transnetyx for genotyping or performed in-house. For in-house genotyping, tissue was incubated overnight at 55°C in One-Step Tail Buffer (100 mM Tris-Cl, pH8.3, 200 mM NaCl, 5 mM EDTA, 1% tergitol, 5 mg/ml ProteinaseK), followed by 1h heat-inactivation at 85°C. PCRs were performed with GoTaq polymerase (Promega).

### EXPERIMENTAL METHODS DETAILS

#### Genomic, transcriptomic, and DNA methylation datasets

We assembled a retrospective multi-institutional medulloblastoma cohort by harmonizing molecular and clinical data from St. Jude and Children’s Oncology Group clinical trials (SJYC07, SJMB03, ACNS0331, and ACNS0332), published multi-omic medulloblastoma cohorts and meta-analyses, and pediatric brain tumor repositories (ClinGen and CBTN). Publicly available molecular data were obtained from primary publications and associated data repositories (GEO: GSE85218, GSE93646, GSE104728). Data from the SJYC07, SJMB03, ACNS0331, and ACNS0332 trials are accessible through interactive St. Jude medulloblastoma data portals. Samples without clinical annotation of patient sex were excluded from further analysis. Across sources, we integrated whole-genome/ exome sequencing, bulk tumor transcriptome profiles (RNA-seq or expression arrays), and DNA methylation arrays. Further, we harmonized sample identifiers, clinical annotations, and molecular subgroup and subtype labels to a common analysis framework.

#### Sex annotation and consistency checks

Patient sex annotations were obtained from clinical metadata provided by originating clinical trials, published cohorts, and repository-derived datasets. Because somatic alterations affecting sex chromosomes, particularly loss of the Y chromosome (LOY) in male tumors, can confound tumor-based sex inference, reported sex was treated as the primary sex annotation. For samples with available matched germline DNA sequencing, molecular sex was validated using germline sequencing data by evaluating relative X and Y chromosome dosage compared to autosomes. Germline X chromosome dosage was used as the primary indicator of biological sex, with Y chromosome dosage serving as a supporting marker in males. Concordance between reported sex and sequencing-based X and Y chromosome dosage was assessed for all samples with available germline genomic data. Samples showing discordance were excluded from downstream sex-stratified analyses.

#### Sample de-duplication via SNP profiling from DNA methylation arrays

To prevent inclusion of duplicate tumors across clinical trials, published cohorts, and repository-derived datasets, we performed sample de-duplication using SNP genotyping probes included within Illumina Infinium DNA methylation arrays (HumanMethylation450K and MethylationEPIC). These arrays contain a set of polymorphic SNP probes designed for sample identity verification and have been widely used for cross-cohort sample matching and detection of sample redundancy in large-scale cancer methylation studies. For each sample profiled on 450K or EPIC arrays, we extracted probe intensities corresponding to SNP probes and constructed a SNP feature matrix. Pairwise sample similarity was assessed by computing all-by-all Pearson correlations between SNP probe intensity vectors. Self-correlations were excluded from downstream analyses. To enable comparison between samples assayed on different array platforms, analyses were restricted to SNP probes shared between the 450K and EPIC arrays, and cross-platform correlation matrices were computed using these common probes. This approach allowed robust identification of duplicate samples across platforms and data sources. Pairs of samples exhibiting a SNP correlation coefficient greater than 0.85 were considered to represent the same biological specimen or technical replicate. In cases where highly correlated samples were identified, a single representative sample was retained based on data completeness (availability of WGS/ WES or RNA-seq/microarray data) and the presence of accompanying clinical annotations, and redundant samples were excluded from downstream analyses.

#### Medulloblastoma subgroup and subtype classification

Medulloblastoma molecular subgroup assignments (WNT, SHH, Group 3, and Group 4) were derived using DNA methylation-based classification where available. High-confidence subgroup annotations from previously published methylation-classified cohorts were used to train a Random Forest classifier for medulloblastoma subgroup prediction. Specifically, samples with established subgroup labels based on methylation profiling were used as a training set, and the trained classifier was applied to predict medulloblastoma subgroup for tumors with available DNA methylation array data. For samples lacking DNA methylation data, medulloblastoma subgroup assignments were obtained from clinical annotations provided by the originating clinical trials or published studies, where subgroup classification had been determined using trial-standard clinical diagnostics. These annotations were used without modification and were not reclassified. Medulloblastoma subtype classification was performed using a two-step approach. First, the Molecular Neuropathology (MNP) version 12.8 methylation classifier was applied to samples within the methylation-classified reference cohort to obtain subtype predictions. These subtype labels were restricted to high-confidence calls and used to define a subtype-labeled training set. A Random Forest medulloblastoma subtype classifier was then trained on methylation array data from this reference set and applied to predict MB subtype for all tumors with available methylation data. Only primary tumor methylation profiles were used for subgroup and subtype prediction. Samples with ambiguous or low-confidence classification results were excluded from subtype-level analyses.

#### Variant calling

Somatic and germline variant calls were obtained using a combination of previously published trial-level variant datasets and uniform reprocessing of raw sequencing data from public repositories. For samples originating from published clinical trials and cohorts with available curated variant calls, we used the variants as reported in the original publications and associated data releases. These variant calls were generated using trial-standard pipelines and quality-control procedures and were incorporated as-is. For samples obtained from ClinGen and the Childhood Brain Tumor Network (CBTN) with available raw sequencing data, we performed standardized variant calling using the nf-core/ nf-sarek Nextflow pipeline (version 3.4.0). For samples with matched tumor–normal pairs, germline variants were called from normal DNA using DeepVariant and GATK HaplotypeCaller, while somatic variants were identified from tumor–normal pairs using Mutect2 and Strelka. For samples lacking matched normal DNA, somatic variant calling was restricted to tumor-only workflows as implemented in nf-sarek. Variant calling was performed on aligned sequencing data generated using pipeline-default best-practice settings. Downstream analyses were restricted to high-confidence variant calls passing caller-specific quality filters.

#### Copy number analysis

Genome-wide copy number alterations were inferred from DNA methylation array data using the conumee R package. Segmental copy number profiles were generated from Illumina Infinium HumanMethylation450K and MethylationEPIC arrays using probe intensity information following background correction and normalization. For each sample, total copy number was estimated by comparing tumor probe intensities to a reference set of normal samples matched for array type. Log ratio-based copy number estimates were calculated across the genome and segmented using circular binary segmentation to identify regions of consistent copy number gain or loss. Segment-level copy number values were used for downstream analyses. Copy number estimates derived from methylation arrays were interpreted as relative copy number states rather than absolute integer copy numbers. Sex chromosomes were analyzed separately from autosomes to account for differences in baseline copy number between males and females. Somatic alterations affecting sex chromosomes, including loss of chromosome Y and alterations in X chromosome dosage, were retained and analyzed as tumor-specific copy number events. Only samples with high-quality methylation data and reliable segmentation profiles were included in copy number analyses. Samples with excessive noise or failed segmentation were excluded from downstream analyses.

#### Expression microarray integration and batch correction

Gene expression microarray data from the Landscape and Cavalli medulloblastoma cohorts were processed and integrated for downstream analyses. Raw expression data were background corrected, normalized, and summarized at the gene level using the Robust Multi-array Average (RMA) algorithm. Probe sets were mapped to gene symbols using platform-specific annotations, and where multiple probe sets mapped to the same gene, expression values were summarized to a single gene-level measurement. Following normalization, expression matrices from the two cohorts were merged based on the intersection of shared genes. To account for systematic technical differences between studies, batch effects attributable to cohort of origin were corrected using the ComBat algorithm implemented in the sva package. Batch correction was performed after normalization and prior to downstream analyses, while preserving biological covariates of interest. Post-correction expression values were used for all downstream gene expression analyses. Samples with missing metadata or evidence of technical artifacts were excluded prior to integration.

#### Differential expression analysis – bulk RNAseq

Differential gene expression analyses were performed using linear modeling and empirical Bayes moderation as implemented in the limma framework. Gene-level expression matrices were modeled using linear models with experimental condition encoded in the design matrix. For each gene, expression levels were fit to the specified design using ordinary least squares regression. Contrasts of interest were defined to compare tumors with loss of inactive X chromosome (LOX) against wild-type (WT) tumors. Contrast-specific model coefficients were estimated using the fitted linear models, and variance estimates were stabilized across genes using empirical Bayes moderation to improve statistical power. Differential expression statistics were extracted for the specified contrast, and genes were ranked based on moderated t-statistics. Statistical significance was assessed using two-sided tests, and multiple hypothesis testing was controlled using the Benjamini–Hochberg false discovery rate (FDR) procedure. All genes passing quality-control filters were included in the analysis.

#### Dissections and flow cytometry

For UBC lineage quantification, *Tbr2-IRES-GFP; LSL-TdTomato* homozygous knock-in male animals were paired with *Lmx1a-Cre* female animals. For FCG experiments, *Sry+* XY male animals were paired with *Tbr2-IRES-GFP* homozygous knock-in female animals. At embryonic day E16.5, cerebellum from individual embryos were dissected into single tubes containing HABG media (Hibernate-A Medium, 2% B27+RA, Glutamax 200nM, 1X Gentamicin). Further processing of the tissue was done using the Papain Dissociation System from Worthington (LK003153). Extracted single cells were prepared for FACS by concentrating to 1×10^6^ cells/ml in FACS buffer (1x PBS, 0.1% BSA). Flow cytometry and cell sorting were performed on a BD FACSAria (Becton Dickinson), gating for live cells with DRAQ7 (Invitrogen, D15106). Data was analyzed with FlowJo or BD FACS-Diva software.

#### Single-cell RNA-sequencing and single-nuclei Multiome-sequencing

For single-cell/nuclei sequencing experiments, FACS-sorted cells were diluted to 20,000 cells in 40 μl FACS buffer. Nuclei were extracted using the 10X Genomics Chromium Nuclei Isolation Kit (1000493), following manufacturer’s instruction. Cells were processed using the 10x Genomics Chromium Single Cell 3’ Reagent Kit (v3.1, PN-1000127), GEM-X Universal 3’ Gene Expression (v4, PN-1000686), and Chromium Next GEM Single Nuclei Multiome ATAC + Gene Expression (PN-1000281), following manufacturer’s instruction. Libraries were sequenced with NextSeq™ 1000/2000 P2 XLEAP-SBSTM Kit (Illumina, 20100987).

#### Single-Cell Analysis

All single-cell and single-nucleus RNA samples were mapped to their corresponding reference genome using Cell Ranger (v8.0.1). Mouse samples were aligned to the GRCm39 prebuilt reference (refdata-gex-GRCm39-2024-A), and organoid samples were aligned to the GRCh38 prebuilt reference (refdata-gex-GRCh38-2024-A), both provided by Cell Ranger. Cells retained by Cell Ranger’s cell-calling algorithm were used for downstream analyses.

Ambient RNA contamination was corrected using CellBender^59^, and cells with a cell probability < 0.9 were removed as compromised. Cells with >10% mitochondrial reads and nuclei with >1% mitochondrial reads were also filtered out. Additionally, cells with gene counts or read counts outside 3 standard deviations above and 1 standard deviation below the sample mean were excluded. Nuclear fraction was estimated using DropletQC^60^, and cells outside 2.5 standard deviations per sample for UMI counts and nuclear fraction were filtered out. Hard thresholds were applied with all cell must have nuclear fraction ≥0.5 for single-nucleus samples and ≥0.1 for single-cell samples, with UMI cutoffs of 2.5 and 3 for single-cell and single-nucleus data, respectively. Ribosomal and mitochondrial genes were removed from downstream analyses to minimize batch effects between single-cell and single-nucleus datasets. Standard Scanpy^61^ workflows were applied for normalization and regression of total counts, pct mitochondrial reads, and cell cycle difference signal. Initial cell annotation was performed by transferring labels from an in-house human prenatal cerebellar atlas using scArches with count matrix, and annotations were then manually curated based on the expression of canonical cell-type markers. Non-neuronal and non-cerebellar cells were removed. After filtering, single-cell and single-nucleus integration was performed using scVI^62^ with 50 latent dimensions. Cell distances were computed in scVI latent space using cosine similarity, and UMAP was generated with min_dist=0.2 and spread=0. Diffusion maps were constructed using Palantir^63^, and pseudotime was estimated using scFates^64^. Single-cell gene set enrichment analysis was performed using Scanpy’s score_genes function. Differential gene scores across pseudotime were assessed by binning cells into 20 equal intervals based on pseudotime. The significance of score differences between male and female mice was evaluated within each bin using a two-sided t-test after aggregating cells from the same sample into pseudobulk profiles.

#### Single-cell multiome analysis

Single-cell RNA and ATAC sequencing data were processed using Cell Ranger ARC v2.0.2 (10x Genomics) and aligned to the mouse reference genome GRCm39 (refdata-cellranger-arc-GRCm39-2024-A) with default parameters. RNA data were filtered and processed as described in the “Single-Cell Analysis” section above. ATAC profiles corresponding to Cell Ranger–identified cells were retained for downstream analyses. After mapping, ATAC data were analyzed using ArchR^65^, with a custom reference constructed from Cell Ranger gene annotations to ensure consistency with mapping reference. Cells with fewer than 1,000 fragments or TSS enrichment < 4 were excluded. Only barcodes present in both RNA and ATAC datasets were kept, and matched RNA expression was added to the ArchR project using addGeneExpressionMatrix. Multimodal neighborhood graphs were constructed in Seurat^66^ using FindMultiModalNeighbors, integrating 50 dimensions from the RNA scVI latent space and 29 dimensions from the ATAC iterative LSI space (LSI dimensions 2–30, excluding the first dimension usually representing sequencing depth). Joint UMAP embeddings were generated using RunUMAP. ATAC peaks were called using MACS2^67^ on ArchR-generated pseudobulk profiles following Leiden^68^ clustering. TF motif activity was quantified using chromVAR^69^ deviation scores via ArchR’s addDeviationsMatrix. Differential TF activity between male and female samples within GC_UBC progenitors and UBC cells was assessed using getMarkerFeatures (Wilcoxon test), adjusting for TSS enrichment and fragment depth. TFs with FDR < 0.05 and absolute deviation score differences > 0.5 were considered significantly active. Differentially active TFs were further assessed by correlating TF gene expression with their corresponding motifs’ accessibility using correlateMatrices, retaining TFs with correlation > 0.5 and adjusted p-values < 0.05 between their TF expression and motif accessibilty. TF motif footprints were computed using getFootprints with Tn5 bias correction and visualized using a 10-bp smoothing window. Target genes were assignedto TFmotifs based on their proximity with the nearest gene using the distanceToNearest function from GenomicRanges^70^. Differential chromatin accessibility and gene expression between male and female samples within GC_UBC progenitors were evaluated using Wilcoxon tests (matrixTests: https://cran.r-project.org/web/packages/matrixTests/index.html), with Benjamini– Hochberg correction. Genes with increased accessibility and expression in males (ATAC activity: adjusted p < 0.05 & M–F > 0 & RNA expression log2FC > 0) were selected to perform overrepresentation analysis using enrichR with GO Biological Process, MSigDB Hallmark, and WikiPathways Mouse databases.

#### Pseudobulk analysis

Single-cell and single-nucleus data were pseudobulked by cell type and sample by aggregating gene counts across all cells within each pseudobulk. Differential expression analysis was performed using PyDESeq^71^. For the sex-matched atlas, cell type and sample origin (single-cell vs single-nucleus) were included as covariates. For the four-core genotype analysis, comparisons between testes_XX vs testes_XY, ovaries_XX vs ovaries_XY, testes_XY vs ovaries_XY, and testes_XX vs ovaries_XX included cell type as a covariate. For comparisons between testes and ovaries in four-core genotype dataset, both cell type and genotype were used as covariates. In the organoid dataset, differential expression was performed across three conditions (control, EST, DHT) using control as the reference. Geneset enrichment analysis was performed using gseapy.prerank, ranking genes by log2FoldChange * -np.log10(padj) with genesets from MSigDB Hallmark 2020 and WikiPathways 2019 Mouse and GO Biological Process 2025. Single-sample GSEA (ssGSEA) was performed using gseapy. ssgsea^42^. Significance testing of ssGSEA scores between two conditions or treatment groups was performed using the ttest_ind function from scipy.stats^72^.

#### Differential Abundance Analysis

Differential abundance analysis in the sex-matched atlas across pseudotime was performed by binning cells into 20 equal intervals based on pseudotime. The abundance of *Mki67*+ cells was compared between male and female mice within each bin using a two-sided t-test after aggregating cells from the same sample into pseudobulk profiles. For the four-core genotype dataset, cell-agnostic differential abundance analysis was conducted using Milo, which clusters transcriptionally similar cells into neighborhoods and tests for differences in neighborhood abundance across conditions. Differences in cell abundance between embryos with ovaries versus testes were assessed across all neighborhoods, using genotype as a covariate.

#### Immunofluorescence (IF) staining

For IF, organoids were fixed in 4% PFA for 30 min, then incubated overnight in 30% sucrose/PBS. Mouse brain samples were fixed in 4% PFA for 2-4h, then incubated overnight in 30% sucrose/PBS. Slides with 10µm-thick sections were blocked for 1 h at room temperature with blocking solution (10% Donkey serum, 0.1% TritonX-100 in PBS). Primary antibodies were diluted in blocking solution and incubated overnight at 4°C. After washing with 0.1% TritonX-100 in PBS for 30 min, secondary antibodies were incubated in blocking solution with DAPI (1 mg/ml) for one hour at room temperature. After washing with 0.1% TritonX-100 in PBS for 30 min, coverslips were mounted on slides. Images were acquired at Zeiss LSM900 Airyscan2, and processed using FIJI or Arrivis Pro (Zeiss). Image quantifications performed with Arrivis Pro Software (Zeiss) using cellpose segmenter. Specificity and sensitivity measurements were performed using the M1 and M2 Colocalization Coefficients.

#### Maintenance of human iPSCs

hiPSC lines (AH017.3^73^ and KOLF2.1J^74^) were maintained in mTeSR+ (StemCell, 100-0276) in Matrigel-coated plates (Corning, 354277) at 37°C, 5% CO2. Once cells were ∼80% confluent, they were split as colonies using ReLeSR (Stem Cell Technologies, 100-0484) for 2 min at 37 °C. Colonies were detached by tapping, and passaged at 1:20 ratio. hiPSCs were cryopreserved as colonies in Bambanker (Nippon Genetics, BB03). Absence of mycoplasma was confirmed at banking via PCR with the PCR Mycoplasma Detection Kit (Abm, ABM-G238), following manufacturer’s instructions. Mycoplasma detection was then performed routinely, and cultures were maintained as mycoplasma free.

#### Generation of transgenic iPSC lines

Human iPSC lines AH017.3 and KOLF2.1J were modified to express a doxycycline-inducible *Cre recombinase* at the *AAVS1* safe locus and engineered to express *a P2A-mNeonGreen-T2A-rtTA-V10* cassette at the 3’ end of the *LMX1A* locus. To generate these reporter lines, hiPSCs were detached using Accumax (Invitrogen, 00-4666-56) for 5 min at 37°C. The Cas9 ribonucleotide protein (RNP) complex was formed by incubating 0.3µl of Cas9 protein (IDT, 1081058) with 22µM sgRNA in Buffer R (Invitrogen, N1025) at room temperature. 2.5×10^5^ cells were electroporated with 1µg of donor plasmid and 1µl of Cas9 RNP complex using the Neon Transfection System (Invitrogen, 1200V, 20ms pulse width, 2 pulses). Cells were plated on Laminin521 (Biolamina, LN521) coated plates in mTeSR+ supplemented with CloneR (StemCell, 05888). After recovery of the cells, single cells/colonies were either selected by puromycin resistance (2 µg/ml for 2 days), or sorted via FACS using BD FACSAria (Becton Dickinson) if a fluorescent marker was present. Genomic DNA was extracted using the DNeasy Blood & Tissue Kit (Qiagen, 69504). Molecular karyotyping was performed using Infinium™ HumanCytoSNP-12 v2.1 Beadchip kits (Illumina), following manufacturer’s instructions. To validate lines, cell line authentication was performed using Multiplex Cell Authentication by Multiplexion GmbH (Heidelberg, Germany, as described^75^. Whole Genome sequencing was performed on Illumina NovaSeq 6000, S4 V1.5 flow cell (Illumina) to confirm the absence of off-targets cuts by the Cas9, and to validate the presence of the KI constructs. KOLF2.1J reporter lines were confirmed as mutation-free. AH017.3 reporter lines underwent a loss-of-heterozygosity of the 17p arm, resulting in a homozygous mutant *TP53* mutation, similar to other reporter lines generated with this hiPSC line^76^. For determination of pluripotency after reporter line generation, human iPSCs were detached using Accumax (Invitrogen, 00-4666-56) for 5 min at 37°C. 5×10^5^ cells were stained with antibody diluted in FACS buffer (2% FBS in DPBS) for 1h at 4°C. After two washes with FACS buffer, fluorescent signal was acquired using a BD LSRFortessa (Becton Dickinson), and analyzed with FlowJo or BD FACS-Diva software.

#### Generation of cerebellar organoids

Organoids were generated following a recently published protocol^43^ with slight modifications. Briefly, iPSCs at 70-80% confluence were washed with PBS and incubated with Accumax for 5 min at 37 °C. The detached iPSCs were then washed and counted before being seeded at a density of 2.4×10^6^ cells per 2 mL of mTESR supplemented with a CEPT cocktail^77^, in one well of an anti-adhesion-treated AggreWell 800 24-well plate (StemCell, 34815). After seeding, the plate was centrifuged at 100g for 3 min. After 24 hours, the 1.5 ml of conditioned media were replaced with 2 ml of fresh mTESR+ without CEPT. For the next two days, 2 mL of conditioned media were replaced with 2 mL of fresh mTESR+. To induce cerebellar differentiation, the spheres were transferred to a 10cm Petri dish containing 10 mL of growth factor-free, chemically defined medium plus insulin (gfCDM): 24% IMDM + GlutaMAX (Gibco, 31980030); 24% Ham’s F-12 (Gibco, 31765035); 5 mg/mL BSA (Sigma-Aldrich, A9576); 1% CD lipid concentrate (Gibco, 11905031); 1% penicillin–streptomycin (Gibco, 15290018); 450 µM monothioglycerol (Sigma-Aldrich, M6145); and 15 µg/mL apo-transferrin (Sigma-Aldrich, T1147), 7 µg/mL insulin (I9278, Sigma), supplemented with 10 µM SB431542 (S1067, Selleckchem), 100 nM LDN-193189 (Adooq Bioscience, A11478) and 1.7 µM CHIR-99021 (Axon Medchem, 1386). From the day of induction, the spheres were cultured on an orbital shaker (Thermo Scientific, 888811102) at 70 rpm. Media changes were performed every other day until day 16. SB431542, LDN193189 and CHIR99021 were added to the medium from days 0 to 10. On days 4 and 6, 100 ng/mL of FGF8b (Thermo Fisher Scientific, 100-25) was added to the medium. On days 6 and 8, the concentration of FGF8b was increased to 300 ng/mL. On day 6, gfCDM was replaced with cerebellar induction medium I (DMEM/F12 + GlutaMAX (Gibco, 31331028), 10% knockout serum replacement (Gibco, 10828028), 0.1 mM beta-mercaptoethanol (Sigma-Aldrich, M6250), 1% penicillin–streptomycin, 15 µg/ml apo-transferrin and 7 µg/ ml insulin). From day 12 to 16, the total medium volume was increased to 15 mL. From day 16 onwards, half-medium changes with cerebellar differentiation medium II (DMEM/ F12 + GlutaMAX, 1% N2 supplement (Gibco, 17502048), 1% B27 minus vitamin A (Gibco, 12587010) and 1% penicillin–streptomycin) were performed every two to three days. From days 30 to 60, the organoids were cultured in a cerebellar differentiation medium (comprising 47% DMEM/F12, 47% Neurobasal-A (Gibco, 10888022), 1% GlutaMAX (Gibco, 35050061), 1% N2 supplement, 1% B27 minus vitamin A, 1% CD lipid concentrate, 1% penicillin/ streptomycin (Gibco, 15140122), 5 µg/ml heparin (Sigma, H3393), 0.25 µg/ml amphotericin B (Gibco, 15290017), 1% growth factor-reduced Matrigel (Corning, 354230) and 100 ng/ml SDF-1 (Gibco, 300-28A) and 50 ng/ml T3 (Merck, T6397). Full medium changes were performed every 7 days.

#### Hormone treatment cerebellar organoids

Five independent batches of organoids were generated as described above, and treated with hormones similar to a published study^78^. At day 16 of differentiation, each batch was split into three plates and treated with ethanol solvent control (96% ethanol, 0.0002 % v/v), DHT (30nM) or estradiol (100nM) were added to the cultures. Media was refreshed every other day and fresh hormones or solvent was added. At day 30 of differentiation, organoids were dissociated with Accumax and flow cytometry/cell sorting was performed on a BD FACSAria (Becton Dickinson). Dead cells were stained with DRAQ7 DROP & GO dye (Invitrogen, D15107) and excluded from downstream analysis. Data was analyzed using FlowJo or BD FACS-Diva software.

